# Terminal selector and subtype selector function across 200 million years of nematode evolution

**DOI:** 10.64898/2026.09.04.749377

**Authors:** Yasmin H. Ramadan, Curtis M. Loer, Hyunsoo Yim, Luke T. Geiger, Daniel M. Merritt, Itai Antoine Toker, Joke Evenblij, Hanh Witte, Steven J. Cook, Ralf J. Sommer, Oliver Hobert

## Abstract

The evolution of brains is subject to investigation in many different animal groups, each offering unique advantages to advance our understanding of the cellular, molecular and regulatory substrates of evolutionary change. Here, we use two nematode species, *C. elegans* and *P. pacificus*, separated by more than 200 million years of evolution to explore how neuronal cell types and the regulatory programs instructing the identity of these cell types have evolved over time. Using gene expression pattern analysis, we compare the differentiation programs of over half of all the nematode’s neuron classes. To explore how the gene regulatory architecture of neuronal differentiation programs evolves, we apply our deep understanding of neuronal differentiation programs, controlled by terminal selectors and subtype selectors in *C. elegans*. Through mutant analysis of orthologous *P. pacificus* regulatory factors, we elucidate patterns of conservation and novelties over such substantial evolutionary distance. We discovered striking similarities in terminal selector expression and activities throughout the nervous system but also observed that terminal selectors can acquire novel sites of expression and distinct regulatory capabilities, manifested by changes in effector gene expression and, hence, neuronal phenotypes. Our mutant analysis argues for a buffering of terminal selector function and for an evolutionary lability of differences in closely related neuronal subtypes. Taken together, our analysis reveals molecular substrates of evolutionary change in nervous systems.

## INTRODUCTION

Within every animal phylum, the overall organization of brains tends to be conserved but the size, cellular typology and organization of cells into functional circuitry are subject to substantial evolutionary change. Comparative anatomical approaches, combined with modern day single cell transcriptomic approaches have resulted in great strides toward a better description of the morphological and cellular substrates of evolutionary change in brains (*1–10*). However, while widely used transcriptomic approaches are valuable in describing conservation and divergences among cell types across evolutionary time spans, they cannot easily address questions that relate to the evolution of gene regulatory networks.

Nematode nervous systems offer a particularly attractive model to explore the evolution of cell types, gene expression programs and gene regulatory networks for a number of different reasons (*11*): First, due to the limited complexity and similar overall organization of nematode nervous systems, as well as conserved cellular lineages over large evolutionary distances, it is possible to homologize cell types based on morphological as well as cell lineage criteria. Second, the regulatory programs that control neuronal differentiation programs are well understood in one particular nematode species, *C. elegans*, and such regulatory programs lend themselves to a functional comparison across different nematode species. These regulatory programs, elucidated through genetic loss of function analysis, are characterized by several features. First, neuron type-specific gene expression programs, and hence, neuronal identity, are driven by terminal selector transcription factors that act in unique, neuron type-specific combinations to coordinately control the expression of terminal identity features of a neuron (*12, 13*). Second, neuron classes that are composed of distinct, but functionally and/or anatomically related subclasses, are controlled by subtype selectors that modulate the ability of a class-specific terminal selector to control the expression of subtype-specific genes (*12, 13*). We have recently shown that across several closely related nematode species within a particular nematode genus, *Caenorhabditis*, the expression of such identity regulators is highly conserved (*14*). However, it is unclear to what extent such conservation of expression extends over much larger evolutionary distances and, even more importantly, to what extent the function of these regulators is conserved.

Armed with a deep knowledge of neuronal differentiation programs in *C. elegans*, we have set out to ask how deeply gene regulatory networks that control neuronal identity are conserved, using as comparison a nematode species that diverged what is now estimated to be around 200 million years ago from *C. elegans*, the diplogastrid *Pristionchus pacificus (15)*. These two nematodes populate distinct ecological niches and show a wide range of behavioral adaptations, including predation and self-recognition (*16, 17*). Our recent EM-based anatomical reconstruction of the main head ganglia of *P. pacificus* revealed homology of neuronal cell types to *C. elegans*, based on neuronal soma positions, neurite projection patterns and synaptic connectivities (*18, 19*). However, to what extent these structurally and lineally homologous cell types have molecularly diverged has been left unclear and whether these homologous neuron classes are generated via similar or diverse regulatory mechanisms is even less well known.

We first analyzed to what extent key molecular identity features of homologous neuron classes are conserved among these two species, focusing on expression analysis of terminal selectors and subtype selectors, as well as key neurotransmitter systems. We examined about half of all neuron classes of these two nematodes, including cholinergic, GABAergic, glutamatergic, monoaminergic and peptidergic neuron classes from all functional categories (i.e. sensory/inter/motor neurons), distributed throughout the entire nervous system, observing broad patterns of conservation of the molecular signature of neuron classes, but also notable differences. Using CRISPR/Cas9 genome engineering in *P. pacificus*, we generated null mutant alleles of select terminal selectors and subtype selectors to explore the conservation of the regulatory architecture of individual differentiation programs. Comparing the resulting mutant phenotypes – from behavior to marker gene expression – we discovered striking patterns of conservation as well as novelties between orthologous *P. pacificus* and *C. elegans* terminal selectors and subtype selectors.

## RESULTS

### Genomic sequence survey of neuron identity regulators

Genetic loss of function studies over the past few decades have revealed a rich set of gene regulatory factors that control neuronal identity in *C. elegans* (*20*). They can be broadly classified into (a) terminal selector-type transcription factors that coordinately control many if not all terminal identity features of a neuron; and (b), subclass selectors that control neuronal subclass identity by promoting or antagonizing the activity of terminal selectors in a subset of neuron class members, resulting in a diversification of a terminal selector-controlled neuron class into distinct subclasses. Given the substantial evolutionary distance of *C. elegans* and *P. pacificus*, and the resulting lack of conservation of many genes (*21, 22*), we first surveyed the fully assembled and well-annotated genome of *P. pacificus* for the presence, absence or duplication of orthologs of *C. elegans* terminal selector-type and subtype selector-type transcription factors whose function was defined by previous genetic analysis in *C. elegans*. These include 40 *C. elegans* homeobox genes, five Zn finger transcription factors, five bHLH factors, four HMG box, two ETS domain, two nuclear hormone receptors and two GATA factors (*14*). Due to the preponderance of homeodomain transcription factors as terminal selectors (*23*), we also examined the *P. pacificus* genome for orthologs of *C. elegans* homeodomain transcription factors that have not been analyzed yet for potential terminal selector gene function.

Using reciprocal BLAST searches, as well as the construction of phylogenetic trees, several themes emerged (**Fig. 1, Table S1**): First, *C. elegans* transcription factors with terminal selector function are generally highly conserved in *P. pacificus* (examples are shown in **Fig. 1A-E**), with the notable exception of the HLH-34 bHLH-PAS protein, which we previously noted to be lost in *P. pacificus* (*18*). Intriguingly, this genomic loss correlates with the loss of the neuron, AVH, whose identity is controlled by HLH-34, through a shift in programmed cell death timing (*18*).

**Figure 1.**
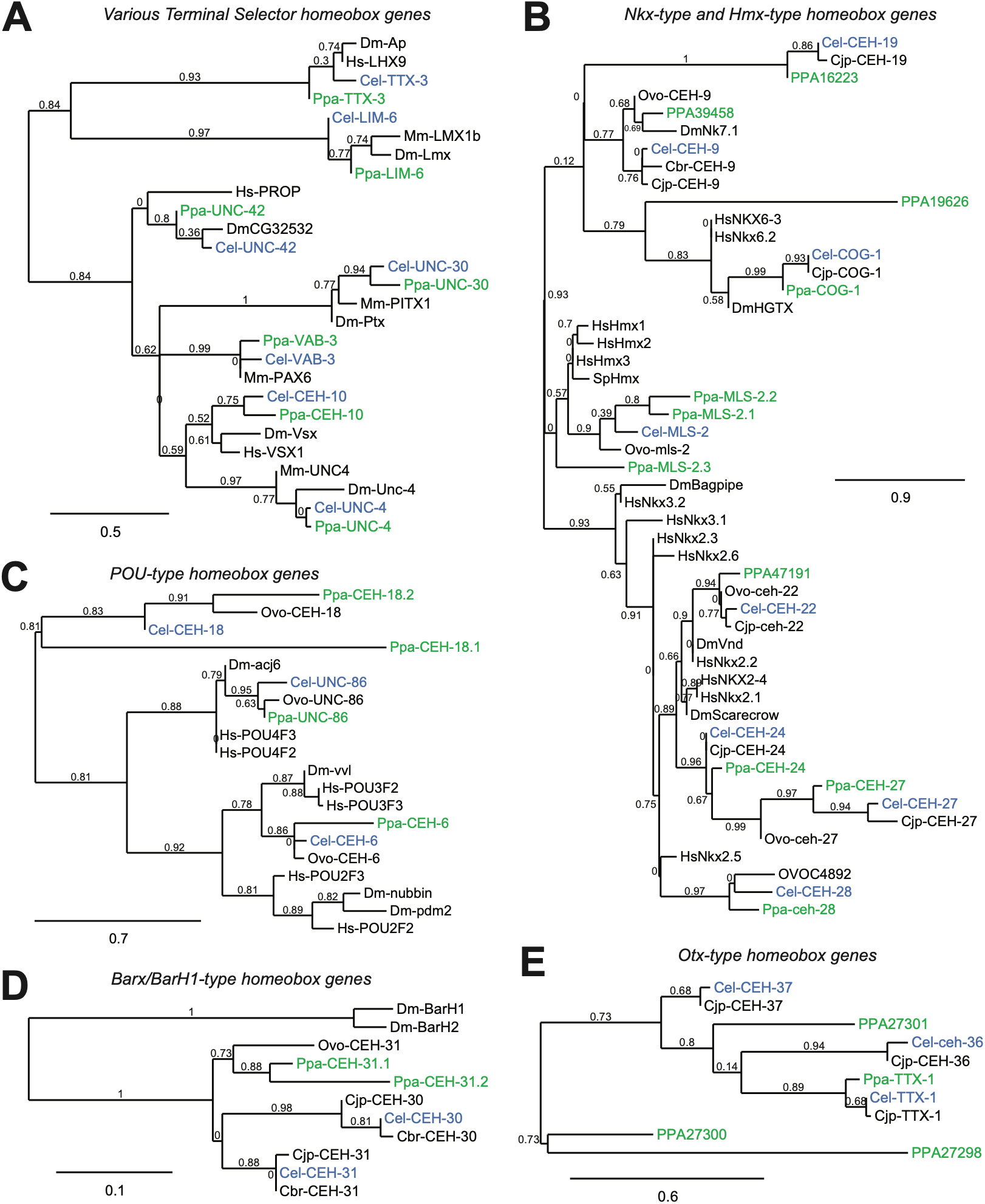
Genomic conservation of homeobox transcription factors and terminal selectors in *P. pacificus* (Ppa) compared to *C. elegans* (Cel). For all trees, generated at the phylogeny.fr server with default values. *P. pacificus (Ppa)* genes are green and *C. elegans (Cel)* orthologs are blue. Branch length scales and node support values are shown. **(A)** Phylogenetic tree showing orthologs of *ttx-3, lim-6, unc-42, unc-30, vab-3, ceh-10*, and *unc-4* in *P. pacificus, C. elegans, D. melanogaster, H. sapiens*, and *M. musculus*. **(B)** Phylogenetic tree of Nkx-type and Hmx-type homeobox genes in *P. pacificus, D. melanogaster, H. sapiens*, and select *Caenorhabditis* species. Note that *mls-2* has three copies in *P. pacificus*. **(C)** Phylogenetic tree of POU-domain-containing homeobox genes *ceh-18, unc-86* and *ceh-6* and their orthologs in *P. pacificus, D. melanogaster, H. sapiens*, and select *Caenorhabditis* species. Note that *ceh-18* has two copies in *P. pacificus*. **(D)** Phylogenetic tree of Barx/BarH1 homeobox gene orthologs in *P. pacificus, D. melanogaster*, and select *Caenorhabditis* species. *ceh-31* duplicated independently in *P. pacificus* and *Caenorhabditis*, resulting in two copies of *ceh-31* in *P. pacificus* and *ceh-30* and *ceh-31* in *C. elegans*, which are both orthologs of the Barx homeobox gene BarH1. **(E)** Phylogenetic tree of Otx-type homeobox genes in *P. pacificus* and select *Caenorhabditis* species. While *ttx-3* has a clear ortholog in *P. pacificus, ceh-36* and *ceh-37* do not, and the *P. pacificus* genome instead contains 2 divergent Otx-type homeobox genes. Additional abbreviations in trees: *C. japonica (Cjp), C. brenneri (Cbr), D. melanogaster (Dm), H. sapiens (Hs), M. musculus (Mm), Onchocerca volvulus (Ovo)* [nematode], *Strongylocentratus purpuratus (Sp)* [sea urchin].

Second, a few terminal selector gene duplications apparent in either *C. elegans* or *P. pacificus* or independent duplications in both (**Fig. 1B-D, Table S2**). Examples include a SIX homeodomain family member, *ceh-34* that duplicated only in *C. elegans* to generate two adjacent paralogs (*ceh-34* + *ceh-33*; only *ceh-34* is expressed in neurons where it acts as a terminal selectors (*24*)). Vice versa, within the NK-superclass, the Hmx ortholog *mls-2*, a terminal selector of several neuron classes in *C. elegans* (*25, 26*), is represented by three copies in *P. pacificus*, two of which are directly adjacent to one another, the third on a different chromosome (**Fig. 1B; Table S2**). A similar duplication event affected a *P. pacificus* member of the POU homeobox family (**Fig. 1C**). The Barx/BarH1 nematode ortholog CEH-31, present in a single copy in more basal nematode classes, duplicated independently in *P. pacificus* and in *C. elegans* (**Fig. 1D; Table S2**). Similarly, of the three Otx-type homeodomains in *C. elegans*, one is deeply conserved in *P. pacificus* (*ttx-1*), while two adjacently located *C. elegans* paralogs, *ceh-36* and *ceh-37*, are represented by three adjacent K50/Otx-type homeobox genes in *P. pacificus*. One of these is most similar to CEH-37, while the other two adjacent genes have significantly diverged (**Fig. 1E**).

All six *C. elegans* members of the CUT homeodomain protein family have been shown to provide critical input into the regulation of pan-neuronal genes (including synaptic vesicle proteins, neuropeptide processing machinery and others) (*27*). Two of these *C. elegans* proteins, CEH-44 and CEH-48, are restricted to all cells of the nervous system while other members are ubiquitously expressed (*27*). In *P. pacificus*, the two panneuronal CUT proteins CEH-44 and CEH-48 are conserved and so is one of the ubiquitously expressed CUT protein, CEH-38 (**Table S1**). A cluster of 3 *C. elegans* CUT homeobox on the X chromosome is not present in *P. pacificus*. A summary analysis of all examined neuronal regulatory factors is provided in **Table S1**.

### Expression patterns analysis of presumptive *P. pacificus* terminal selectors and neurotransmitter identity genes

In considering how best to study selector gene expression in the *P. pacificus* nervous system, we considered two issues: 1) transgenic promoter fusion reporters have a long-standing history in *C. elegans* of revealing only partially correct expression patterns due to lack of *cis-*regulatory elements and 2) in contrast to the insertion of small epitopes, insertion of larger sequences such as fluorophore genes into genomic loci by CRISPR/Cas9 genome engineering has not yet been possible in *P. pacificus*. Therefore, we used primarily two techniques for reliably assessing expression patterns: (1) we tagged endogenous *P. pacificus* loci with FLAG epitopes using CRISPR/Cas9 for some genes (**Supp. Fig. S1**) and (2) we detected endogenous gene transcripts using hybridization chain reaction for in situ detection of mRNA (HCR RNA FISH, in short, HCR) for all other genes, including those that we epitope tagged. For one gene *(unc-86)*, we also examined staining with an anti-*Cel-*UNC-86 antiserum. In total, we analyzed the expression of 31 *P. pacificus* orthologs of *C. elegans* terminal and subtype selector genes with these approaches. HCR permitted us to perform double- or triple stains to assess combinatorial expression of putative terminal selectors.

In parallel to analyzing terminal and subtype selector genes, we also examined the expression of neurotransmitter and neuropeptide genes to probe the extent of conservation of neuronal identity features. These signaling features are archetypical targets of terminal selectors (*12*). In total, our analysis probed the identity of 57 functionally diverse neuron classes, distributed throughout all regions of the central, peripheral and enteric nervous system of *P. pacificus* (**Table 1**), covering about half the entire nervous system of both nematode species.

**Table 1.** Summary of molecular cell typology. Bolded genes and shaded neuron class rows indicate differences to *C. elegans*.

| cell | Transcription factor(s) | neurotransmitter/<br>neuropeptide examined | additional marker<br>examined |
| --- | --- | --- | --- |
| **Head sensory neurons** |  |  |  |
| IL2 | *unc-86* + *cfi-1* + *sox-2.1*<br>**no *unc-39* in IL2L/R** | *unc-17* (ACh) | *klp-6*, *egas-1-9*, *degl-1*<br>**no *flp-14*** |
| URA | *unc-86* + *sox-2.1* + *vab-3*, | *unc-17* (ACh) |  |
| URB | *unc-86* + *sox-2.1* + *vab-3* + *ceh-31.2* | *unc-17* (ACh) |  |
| OLQ, OLL | *vab-3* + *ceh-32* (OLL only) | **no *eat-4* (Glu) – OLQ** | ***ocr-4* (new in OLL)** |
| URY | *unc-86* |  |  |
| ASH | *unc-42* | *eat-4* (Glu) | *osm-6* |
| AFD | *ceh-14* + *ttx-1* + ***che-1*** | *eat-4* (Glu) |  |
| ASEL/R | *che-1* + ***lim-6* (not asymmetric)** | *eat-4* (Glu) |  |
| ASK | *mls-2.2*, *mls-2.3*, **no *ttx-3*** | *eat-4* (Glu) |  |
| ASI | **no *unc-3* (Ab)** |  |  |
| ASG | **no *unc-30* (Ab)** |  |  |
| AUA | *hlh-17* | *eat-4* (Glu) |  |
| **Head interneurons** |  |  |  |
| AIA | *ttx-3* + *unc-39* | *unc-17* (ACh) |  |
| AIY | *ttx-3* + *ceh-10* | *unc-17* (ACh) |  |
| AIZ | *unc-86* + *ceh-43* | *eat-4* (Glu) |  |
| AIM | *unc-86* + *ceh-14*, **no *mls-2*** | *nlp-70*, *nlp-73*, *unc-17* (ACh) in males,<br>**no *eat-4* in hermaphrodites,**<br>**no 5HT-IR, no *mod-5*;** |  |
| ALA | *ceh-14* | *flp-13* |  |
| Command<br>interneurons | *unc-42* + *unc-3* | *unc-17* (ACh) |  |
| AVF | *unc-4* | *snf-11* |  |
| AVJ | *unc-30* + *mls-2.3* |  |  |
| AVK | *unc-42* | *flp-1* (no *eat-4*, *unc-17*, *unc-25*, i.e.<br>remains peptidergic) |  |
| RIC | *hlh-13* | *tbh-1*, *tdc-1* |  |
| RIG | *lim-6* | *eat-4* |  |
| RIH | *unc-86* | *nlp-71*, 5HT, **no *mod-5*** |  |
| RIP | *unc-86* + *ttx-1* | **new 5HT-IR, new *tph-1***<br>**new *unc-25* (but no anti-GABA),**<br>**no *unc-17*, no *nlp-73*** |  |
| RMG | *unc-86* | *nlp-56*, *flp-14* (no *unc-17*),<br>**new *eat-4* asymmetry (RMGR only)** |  |
| **Body mechanosensory neurons** |  |  |  |
| Touch neurons<br>(ALM, PLM,<br>AVM, PVM) | *unc-86* + *mec-3* | *eat-4* (Glu) | *mec-18* |
| all dopaminergic<br>neurons<br>(CEP/ADE/PDE) | *ast-1* + *ceh-43* | *cat-2* (DA) |  |
| **Midbody** |  |  |  |
| HSN | *unc-86* + *ast-1*, **no *hlh-3*** | *unc-17*, no *eat-4* (Glu), **no *tph-1*** |  |
| CAN | *ceh-10* + *ceh-43* | *nrps-1*, no *egl-3*, **no *ceh-48*** |  |
| SDQ | *ceh-43* + *ceh-31.2*, no *unc-86* | *unc-17* (ACh) |  |
| BDU | *unc-86* + *ceh-43* + *ceh-14* + *ceh-31.2* | **new *unc-17* (ACh)** |  |
| **Ventral nerve cord MNs** |  |  |  |
| DA | *unc-3* + *unc-4* | *unc-17* (ACh) | *madd-4* |
| DB | *unc-3* + *vab-7* + *ceh-12*; DB2: *hlh-17* | *unc-17* (ACh), **no *acr-5*** | *madd-4* |
| VA | *unc-3 + unc-4 + bnc-1* | *unc-17 (ACh)* | *madd-4* |
| VB | *unc-3 + ceh-12 + bnc-1; VB2: hlh-17* | *unc-17 (ACh), no acr-5* | *madd-4* |
| AS | *unc-3 + unc-55 + mab-9* | *unc-17 (ACh)* | *madd-4* |
| VC | *unc-4 + vab-7, no hlh-3* | *unc-17 (ACh) + 5HT* | *madd-4* |
| DD | *unc-30, no elt-1.1 or elt-1.2* | *unc-25 + unc-46 + unc-47,*<br>*no flp-13, no oig-1, no acr-14* | *ilys-4, pde-4, madd-4* |
| VD | *unc-30 + unc-55, no elt-1.1 or elt-1.2* | *unc-25 + unc-46 + unc-47,*<br>*no oig-1, no acr-14* | *ilys-4, pde-4, madd-4* |
| **Tail inter- and motoneurons** |  |  |  |
| PVP | *unc-3 + unc-30* | *unc-17 (ACh)* |  |
| DVB | *lim-6* | *GABA-Ab, unc-25, unc-47* |  |
| PVR | *ceh-31.1 + ceh-32.2* | *eat-4* |  |
| **GABAergic head neurons** |  |  |  |
| RME | *lim-6 (RMEL/R only)* | *GABA-Ab, unc-25, unc-47* |  |
| AVL | *lim-6* | *GABA-Ab, unc-25, unc-47* |  |
| RIS | *lim-6* | *GABA-Ab, unc-25, unc-47* |  |
| **Enteric neurons** |  |  |  |
| NSM | *unc-86 + ttx-3* | *5HT-IR, tph-1* |  |
| **Male-specific neurons** |  |  |  |
| MCM | *unc-42* |  |  |
| CEM | *unc-86* |  |  |
| **Glia** |  |  |  |
| GLR glia | *unc-30 + let-381* | *snf-11* |  |

We display first the antibody staining results with four candidate Ppa terminal selectors (UNC-86, UNC-42, UNC-3, UNC-30)(**Fig. 2**, summarized in **Supp. Fig. S2**) and then expand these results with additional HCR-based marker analysis (**Fig. 3** to **Fig. 6**), followed by mutant analysis (**Fig. 7** to **Fig. 11**). Apart from the initial antibody staining figures (**Fig. 2**), the description of results is, however, not organized by individual genes but rather by nervous system regions and cell types. The results of the fate marker analysis are summarized in **Table 1**.

**Figure 2.**
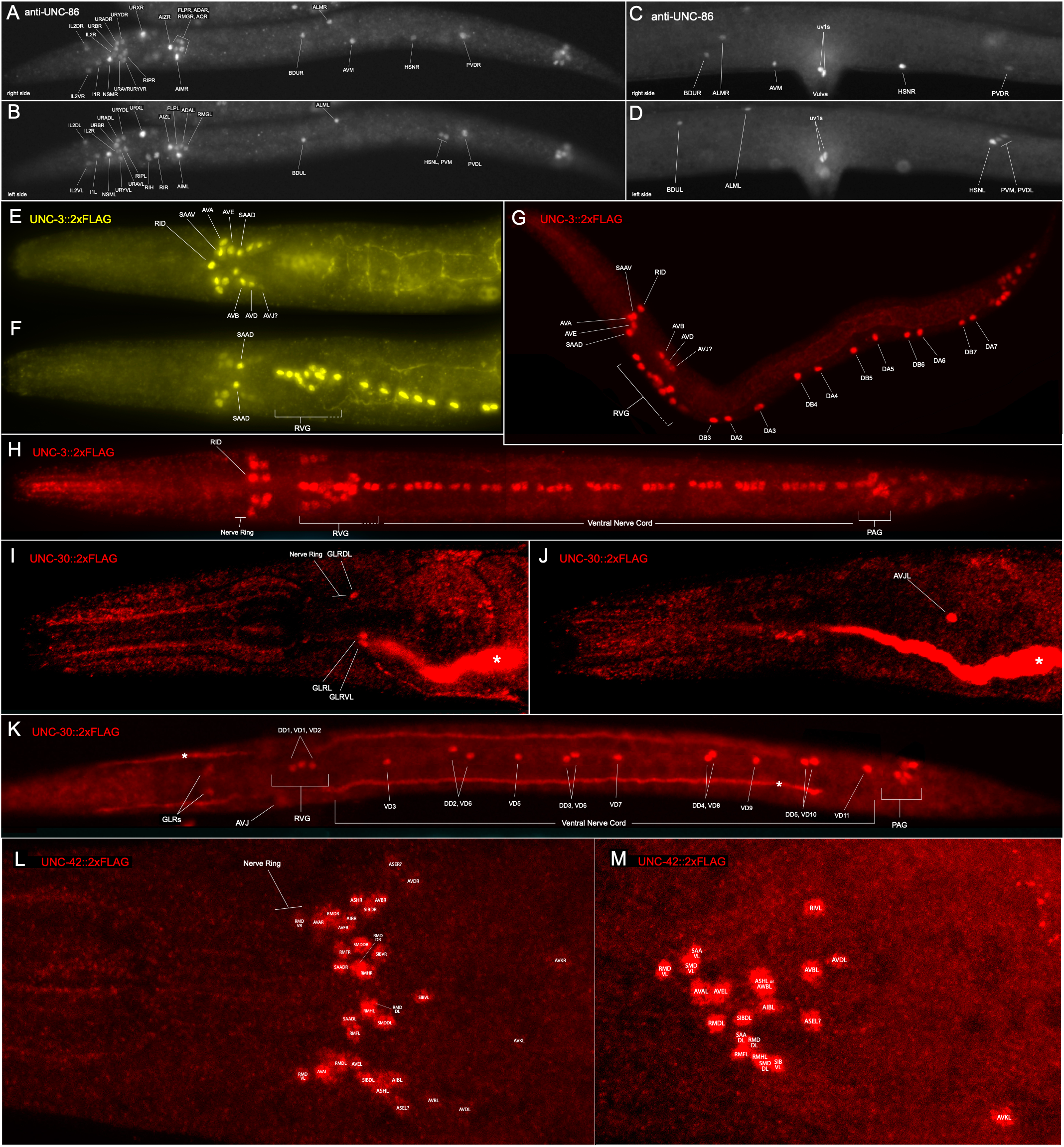
Expression patterns of candidate terminal selectors UNC-86, UNC-3, UNC-30 and UNC-42 by antibody staining reveal conservation of *C. elegans* expression patterns in *P. pacificus*. All cell identifications indicated here are best guesses based on cell body position and conservation relative to *C. elegans*, and are, in part, also corroborated by HCR staining with additional markers, as shown in the ensuing figures. **(A – D)** Anti-*Cel*-UNC-86 staining in *P. pacificus*. (A) Right side of entire larva demonstrating expression in neuronal nuclei like those seen in *C. elegans*; nuclei in the head and body are identified as indicated. (B) Left side of same larva as in A. The ventral midline unpaired head neurons RIH and RIR are also visible in this focal plane. (C) Right side, central body region of adult hermaphrodite showing body wall neuronal nuclei (same as in the larva) and a pair of vulva-associated uv1 cells seen only in the adult. Note that HSNs are positioned asymmetrically on the anterior-posterior axis, with HSNR closer to the vulva. (D) Left side of same worm as in C. HSNL is located further posterior than HSNR, clustered with two other UNC-86-expressing nuclei, PVDL and PVM. **(E – H)** Anti-FLAG antibody staining on *P. pacificus* strain, *unc-3(tu1823[unc-3::2xFLAG])* in which the endogenous *unc-3* locus has been tagged with a 2xFLAG epitope tag. (E) Late larva head and anterior body, dorsal & lateral focal planes; nuclei are identified as indicated. Some identifications (?) are uncertain. In adults and late larvae, staining of the nucleus marked ‘?’ (likely AVJ, possibly AIZ) is either faint, as seen here, or absent. (F) Ventral focal planes of the same larva as in E, showing the ventral-most head nuclei (SAADs), RVG nuclei and the anterior portion of the ventral nerve cord (VNC). (G) Lateral view of recently hatched larva, prior to P.na divisions generating postembryonic UNC-3-positive neuronal nuclei of the VNC. Nuclei in the head are the same as observed in later larvae and adults (except as noted above). Embryonically generated VNC nuclei belonging to cholinergic motor neurons are identified as indicated and are the same as seen in *C. elegans*. (H) Larva showing complete set of UNC-3-expressing cholinergic motor neuron nuclei in the VNC; montage of a few focal planes, mostly ventral, to better show all VNC nuclei. Location of the nerve ring (NR) is indicated to show that all UNC-3-expressing are posterior to the NR, as seen in *C. elegans*. (I) **(I – K)** Anti-FLAG antibody staining on *P. pacificus* strain, *unc-30(tu2075[unc-30::2xFLAG])* in which the endogenous *unc-30* locus has been tagged with a 2xFLAG epitope tag. * - indicates canal cell staining seen with this anti-FLAG antibody in wildtype worms. (I, J) Head of adult hermaphrodite showing head neuron and glia staining, lateral views (ventral down). (I) Midline focal planes showing GLR glial nuclei adjacent or slightly overlapping the nerve ring. (J) More lateral focal plane showing a single UNC-30-expressing nucleus in the posterior head, AVJ on the left side. (K) Larva showing UNC-30-expressing GABAergic motor neuron nuclei in the RVG and VNC as indicated; head nuclei (seen in I, J) are also indicated but are largely out of focus. **(L, M)** Anti-FLAG antibody staining on *P. pacificus* strain, *unc-42(tu1824[unc-42::2xFLAG])* in which the endogenous *unc-42* locus has been tagged with a 2xFLAG epitope tag. Closeups on head region. A few nuclei with uncertain identification include a ‘?’ All expressing nuclei are located posterior to the nerve ring; no nuclei in the VNC, body or tail express UNC-42. (L) Ventral view showing bilaterally symmetric UNC-42-expressing nuclei More dorsal stained nuclei are not seen in these focal planes, although this ventral view shows 17 of ~20 likely UNC-42-positive nuclei; (R,S) (M) Left lateral view closeup, showing 20 stained nuclei on one side of the head. Likely identifications as indicated.

**Figure 3.**
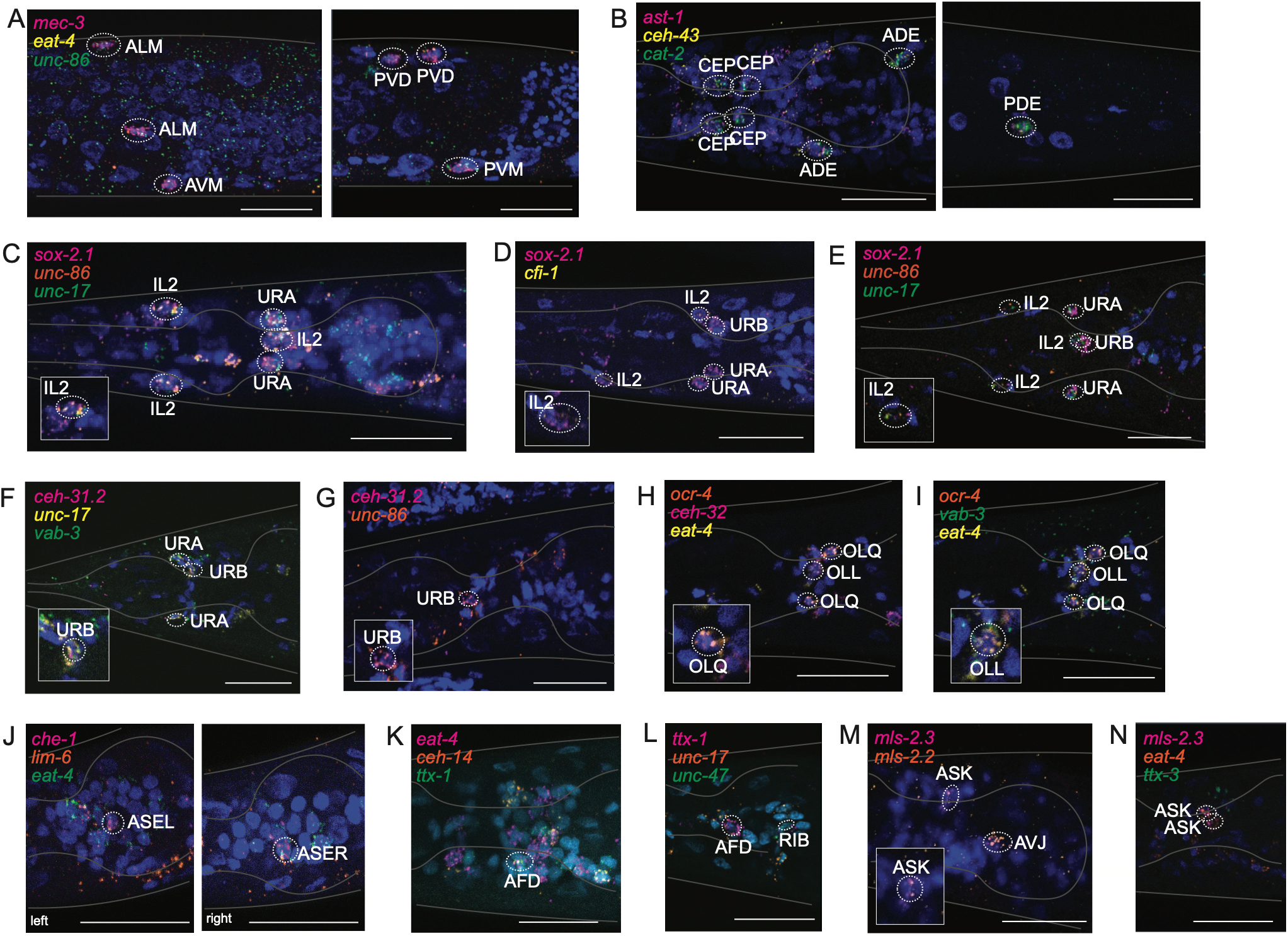
Expression patterns of candidate identity markers in sensory neurons. HCR RNA-FISH staining of various genes as indicated in each panel. Probe details are found in Table S5. Anterior is to the left in all images. In several cases a higher magnification version of parts of the same panel are shown in an inset. **(A,B)** Different types of touch receptor neurons, the glutamatergic light touch receptor neurons (A) and the dopaminergic neurons (B), show conserved molecular signatures of regulatory factors and terminal effector genes. (**C-G)** Cholinergic anterior ganglion neurons (IL2, URA, URB) show conserved molecular signatures. Expression of *ceh-31* paralogs in URA could not be unambiguously determined. (**H,I**) Glutamatergic sensory neurons in the anterior ganglion show similar expression of putative terminal selectors, but changes in the subtype specificity of some effector genes (*eat-4* and *ocr-4)*. Both panels show the same animal, but show staining with different probe sets. **(J-M)** Conservation and novelties of terminal selector ortholog expression in amphid sensory neurons. Expression of *mls-2* orthologs, conserved in ASK, are also conserved in the AVJ interneurons (but no in the AIM interneurons)

**Figure 4.**
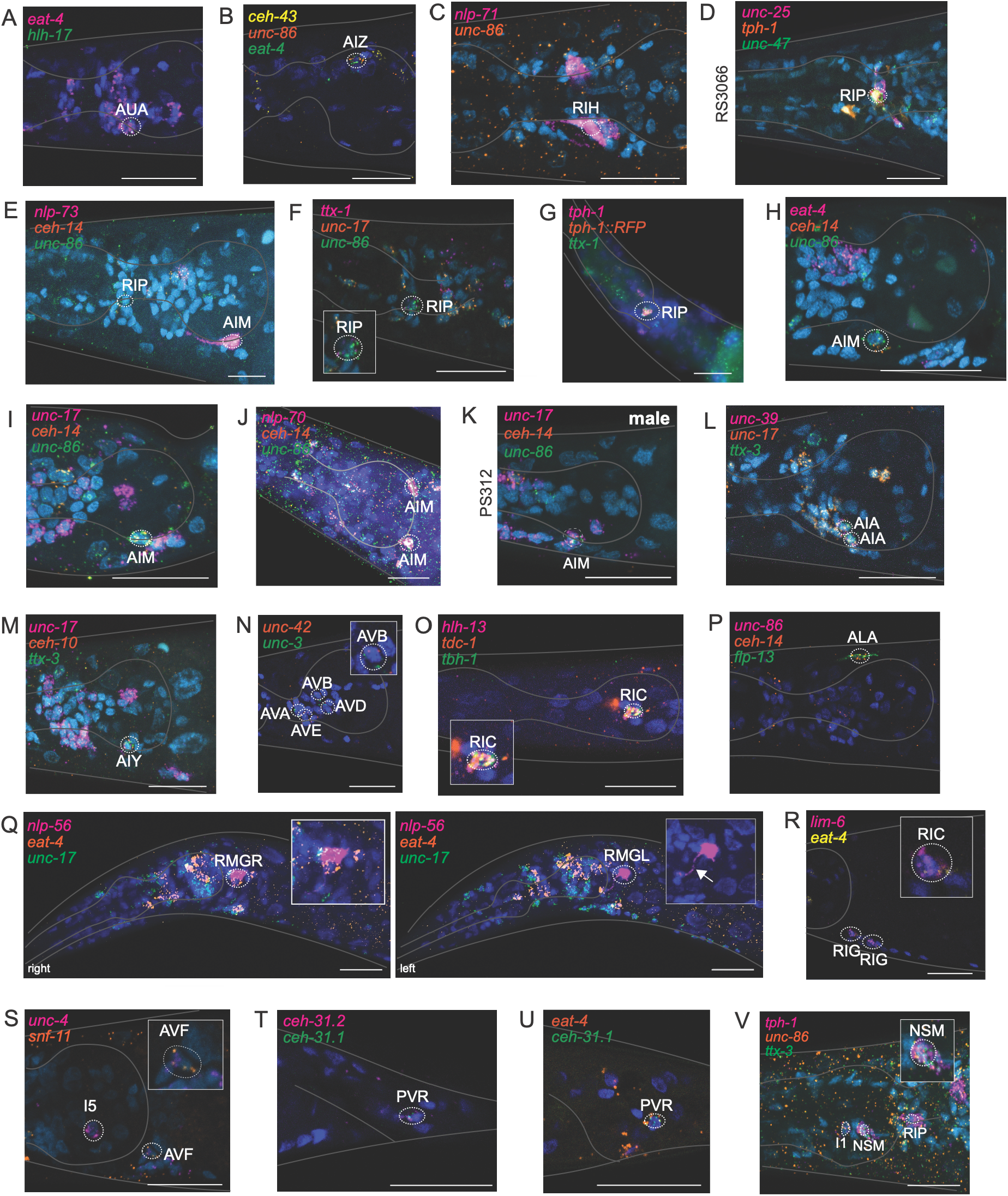
Expression patterns of candidate identity markers in head inter- and motorneurons. HCR RNA-FISH staining of various genes as indicated in each panel. Probe details are found in Table S5. (**A**) Glutamatergic AUA neurons, an unusal neuron class with a shortened dendrite in some, but not all nematode species, shows similar neurotransmitter identity and preserve expression of the AUA terminal selector, *hlh-17*. (**B,C**) *unc-86-*dependent AIZ and RIH interneurons show a similar molecular profile in *P. pacificus*. (**D-G**) The RIP neuron class, the key connector between the central and pharyngeal nervous system, shows a preserved regulatory signature *(C. elegans* terminal selectors *unc-86* and *ttx-1*) but divergent neurotransmitter features. *tph-1::rfp* indicates that staining was done on an transgenic line expressing *tph-1::rfp* (in serotonergic neurons, including RIP, as previously reported (*37*)). The array-expressing RIP neuron is visualized with HCR probes against RFP. (**H-K**) The AIM neuron shows similarity and differences in molecular profiles (*unc-86, ceh-14*, but not *mls-2*) are conserved. See panel E for conservation of *Ppa-unc-86* in AIM. **(L)** *unc-17/VAChT* is expressed in AIM in male animals, as it is in *C. elegans*. **(M, N)** The cholinergic interneurons AIY and AIA, both requiring *ttx-3* as terminal selector, show conservation in neurotransmitter identity and terminal selector expression (*ttx-3-*dependent AIA and AIY and cofactors *ceh-10* in AIY and *unc-39* in AIA). **(O)** Command interneurons show a conserved overlap of *Ppa-unc-3* and *Ppa-unc-42* expression, co-terminal selectors in *C. elegans*. **(P)** Co-expression of the biosynthetic enzymes for octopamine, *tbh-1* and *tdc-1*, as well as their regulator, *hlh-13*, indicates that the RIC interneuron class is conserved in *P. pacificus*. **(Q,R)** ALA and RMG neurons preserve neuropeptide (*flp-13* in ALA and *nlp-56* in RMG) and terminal selector expression in *P. pacificus* (*ceh-14* in ALA and *unc-86* in RMG) is preserved. Note that *nlp-56* signals are so abundant that axon morphology can be observed (inset with white arrow) and that one of the two RMG neurons also display *eat-4/VGluT* expression, an asymmetric neurotransmitter identity observed nowhere else in the nervous system. **(S,T)** Two interneurons in the retrovesicular ganglion, the glutamatergic RIG neuron and the GABA reuptake neuron AVH show conserved molecular profiles in *P. pacificus*. Note that the regulatory of AVF identity, *unc-4*, also preserves its I5 pharyngeal interneuron expression in *P. pacificus*. **(U,V)** The glutamatergic PVR tail interneuron maintains neurotransmitter identity and expresses both *P. pacificus* copies of its *C. elegans* regulator *ceh-31*.

**Figure 5.**
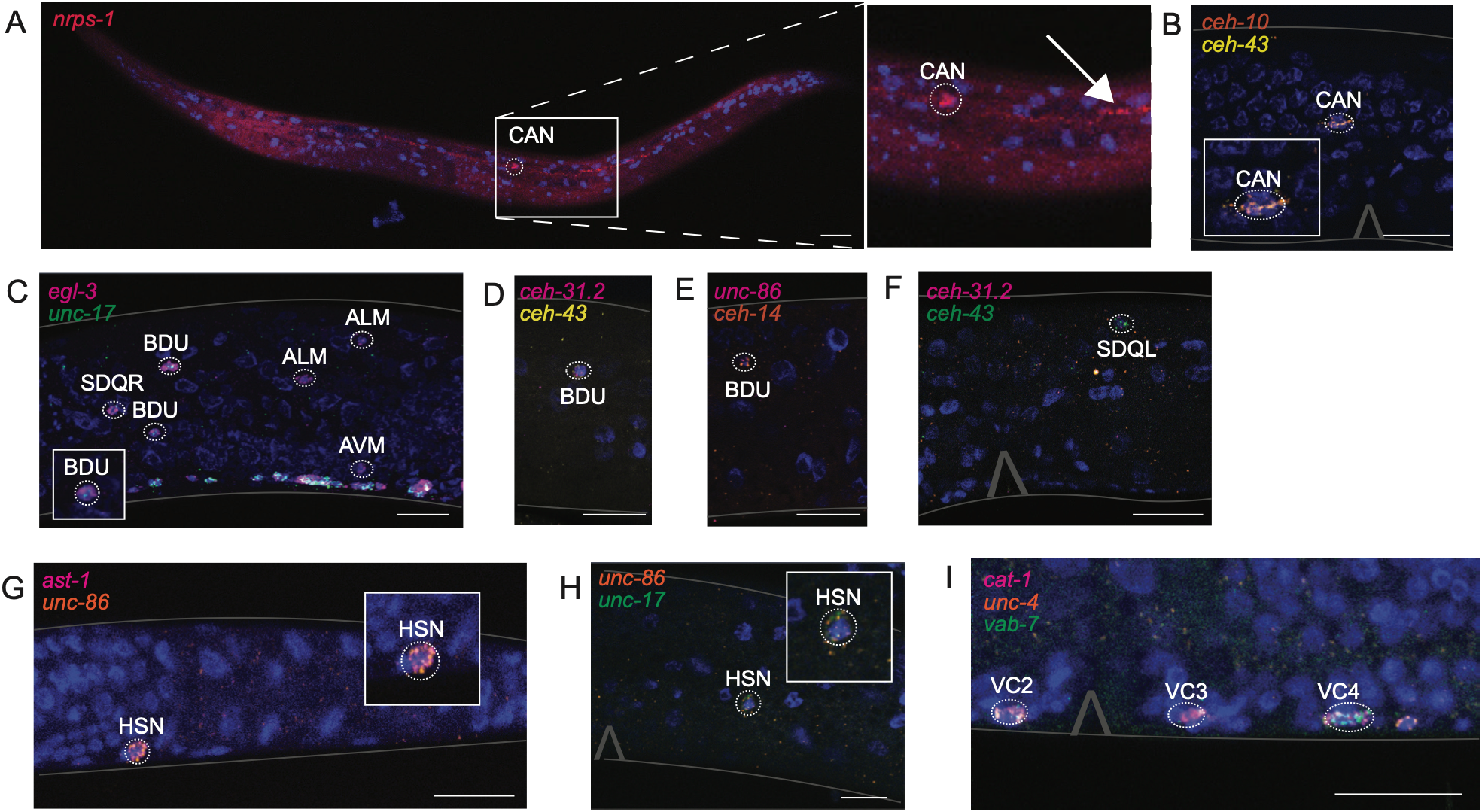
Expression patterns of candidate identity markers in midbody neurons. HCR RNA-FISH staining of various genes as indicated in each panel. Probe details are found in Table S5. (**A,B**) The CAN midbody neurons preserves expression of the nemamides-synthesizing *nrps-1* gene (its expression is so strong that its axon is visible; indicated in blow-up), as well as expression of regulatory factors controlling CAN function and development (*ceh-10, ceh-43*). **(C-F)** Expression of the panneuronal *egl-3* gene, involved in neuropeptide processing, shows overall preservation of midbody neurons, including SDQ and BDU, which also preserve expression of regulatory factor in *P. pacificus*. Note that BDU shows a cholinergic phenotype (*unc-17/VAChT* expression) that it does not show in *C. elegans*. **(G-I)** Putative vulval muscle-innervating HSN and VC neurons show similarities and divergences. HSN expression of *Ppa-unc-86* is preserved, *Ppa-ast-1* expression is very low and serotonergic identity is not conserved (*37*). Expression of VC features *Ppa-unc-4* and *Ppa-vab-7* is preserved; note that these neurons become robustly serotonergic in *P. pacificus* (*37*).

### Peripheral and central mechanosensory neurons show conserved terminal selector gene expression

UNC-86/POU-IV-type homeobox genes control the differentiation program of peripheral sensory neurons across animal phylogeny (*28*). In *C. elegans*, the UNC-86 protein is expressed and required for the function of mechanosensory neurons located in the head, tail and periphery of the animal (FLP, ALM, AVM, PVD, PLM, PVM)(*29, 30*). We identified expression in the same cells in *P. pacificus* via anti-Cel-UNC-86 antibody staining and *unc-86* transcript detection by HCR (**Fig. 2A, B, 3A**). In all these mechanosensory neurons, *C. elegans* UNC-86 is known to cooperate with the MEC-3 LIM homeodomain protein to specify their identities (*31, 32*). We observed co-expression of *Ppa-unc-86* and *Ppa-mec-3* transcripts by HCR (**Fig. 3A**), suggesting that the *unc-86* / *mec-3* combinatorial code is conserved in *P. pacificus* (see below for mutant phenotypes). We found that the glutamatergic identity of the touch receptors neurons, assessed by *eat-4/VGluT* expression (*33*) also to be conserved in *P. pacificus* with the sole exception that, unlike in *C. elegans*, the *P. pacificus* PVM is positive for *eat-4/VGluT* (**Fig.3A**). A complete conservation of expression is also observed for *mec-18* (**Fig. 3A**), one of the many UNC-86/MEC-3-dependent and touch neuron-restricted *mec* genes required for mechanosensory function (*34*).

The total set of eight dopaminergic neurons in *C. elegans*, all mechanosensory neurons as well (*35*), falls into 3 classes, located in the periphery (PDE) as well as head ganglia (ADE, CEP) (*36*). Histochemical staining, plus HCR against dopamine synthesis and transport components show that these neurons are dopaminergic in *P. pacificus* as well (**Fig. 3B**)(*37, 38*). In *C. elegans*, these neurons are defined by a combination of identity controlling terminal selectors, including the ETS domain transcription factor AST-1 and the Distalless/Dlx ortholog CEH-43 (*39, 40*). Through double HCR staining of *Ppa-ast-1* and *Ppa-ceh-43*, as well as dopamine synthesis genes, we find this unique combinatorial code to be preserved in *P. pacificus* (**Fig. 3B**).

### Anterior ganglion neurons show patterns of conservation and divergence

*C. elegans* UNC-86 is also expressed in head neuron classes anterior to the nerve ring, including chemo- and mechanosensory inner labial IL2 sensory neurons, and putative multimodal neurons URA, URB and URY that also send dendritic processes toward the tip of the nose (*30*). The number and position of neurons that express *unc-86* (based on HCR and antibody staining) is similar between *C. elegans* and *P. pacificus* (**Fig. 2A,B**). To further probe the similarities between *C. elegans* and *P. pacificus* anterior ganglion neurons, we examined additional markers. In *C. elegans*, UNC-86 cooperates with HMG-box SOX-2, Pax6/Eyeless ortholog VAB-3 and ARID-type CFI-1 transcription factors to specify the proper identity of the IL2, URA and URB neurons (*26, 41–43*). We find that *Ppa-unc-86, Ppa-vab-3* and a *Ppa-sox-2.1* ortholog are also co-expressed in IL2, URA and URB neurons in *P. pacificus* (**Fig. 2A,B, Fig. 3C-G**).

In *C. elegans*, a nematode BarH/Barx-type homeobox gene *ceh-31* is expressed in both URA and URB neurons (*26*), but not in the IL2s. Of two *P. pacificus ceh-31* orthologs (*ceh-31.1* and *ceh-31.2*), *ceh-31.2* is expressed in URB (**Fig. 3F,G;** its expression in URA is not clear). The ARID-type transcription factor *cfi-1/ARID* co-operates with *unc-86* to specify the *C. elegans* IL2 neurons (*41, 42*) and we find that in *Ppa-cfi-1* is co-expressed with *unc-86* in *Ppa* IL2 neurons as well (**Fig. 3D**). The cholinergic identity of *C. elegans* IL2 neurons requires *unc-86* and *cfi-1* function (*41*). *P. pacificus* IL2 also use acetylcholine, as assessed by cholinergic marker protein and transcript expression (**Fig. 3C**).

In *C. elegans*, the six radially symmetric IL2 neurons can be subdivided into two different subtypes, the lateral IL2 pair and the dorsoventral IL2 pairs, based on gene expression, as well as altered dendritic branching patterns elaborated in the dauer stage (*44–46*). This diversification is controlled by the *C. elegans unc-39* SIX-type homeobox gene, which is expressed only in the lateral IL2s to promote lateral identity and inhibit dorsoventral identity (*45*). Strikingly, expression of *Ppa-unc-39*, while conserved in the AIA interneuron class (described below in **Fig. 4**), is absent in lateral IL2 neurons (**Fig. 7I,J**; data shown in a later figure as part of our mutant analysis described later). Expression of IL2 subtype-specific *C. elegans unc-39* target genes show corresponding differences in *P. pacificus*. In *C. elegans, degl-1* (a degenerin channel-like gene) is expressed in the IL2 D/V pairs, whereas *degl-2* is expressed in the IL2 lateral pair (*45, 46*). In *P. pacificus*, there is a single *degl* ortholog, which we term *degl-1*, that is expressed in all six IL2 neurons (**Fig. 7 I,J**). The FMRF-like peptide gene *flp-14* is selectively expressed in *C. elegans* lateral IL2s (*45*), but its *P. pacificus* ortholog is not expressed in the IL2s (**Fig. 7 I,J**). Three of four *C. elegans egas* genes, encoding EGF-domain-containing ASIC-like channel proteins, show subtype-specific expression (*egas-1* and *egas-3* in dorsoventral IL2s, *egas-4* in lateral IL2s) (*45, 46*). In *P. pacificus*, there are 9 *egas* genes with more modest subtype-specific expression: although all *egas* genes are expressed in all IL2 neurons, two genes, *egas-1* and *egas-2*, are more strongly expressed in the dorsoventral IL2 pair compared to the lateral pairs (**Fig. 7 I,J**). We hypothesize that the absence of *unc-39* expression in the IL2 lateral neuron subtype may contribute to the much less pronounced subtype diversification of IL2 neurons in *P. pacificus*.

**Figure 6.**
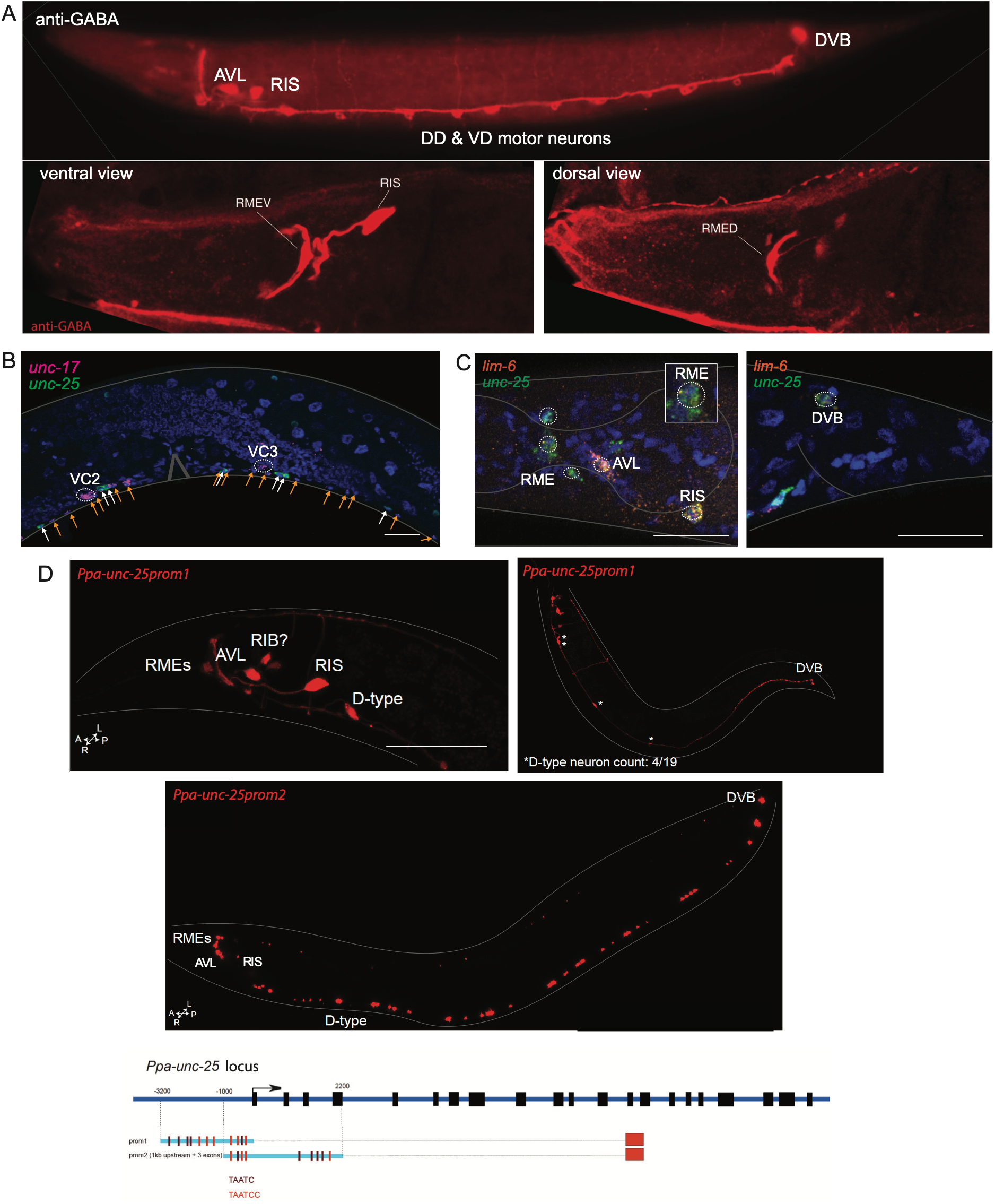
Conservation of GABAergic neurons in *P. pacificus*. **(A)** Anti-GABA staining shows the same complement of main GABAergic neurons in *Ppa* compared to *Cel*, with the exception of staining of the RIB neuron pair which is weak and ambiguous. Other weakly GABA-positive neurons include outer labial neurons (not seen in this preparation). **(B)** HCR RNA-FISH staining of showing interdigitation of GABAergic (*unc-25/GAD*) and cholinergic (*unc-17/AchT)* motorneurons in the *P. pacificus* ventral nerve cord, mirroring the patterns observed in *C. elegans*. **(C)** The *Ppa* ortholog of the GABAergic terminal selector *Cel-lim-6* shows the same expression pattern in *P. pacificus* (RIS, AVL, DVB neurons), marked with *unc-25/GAD*. **(D)** Two transgenic reporter lines, indicated at the bottom of the panel, containing parts of the *unc-25/GAD* locus, fused to RFP, recapitulate the major GABA neuron classes also observed with *unc-25/GAD* HCR. One of the two reporters (prom1), contains fewer predicted UNC-30 binding sites, which may explain the less robust expression in D-type motor neurons.

**Figure 7.**
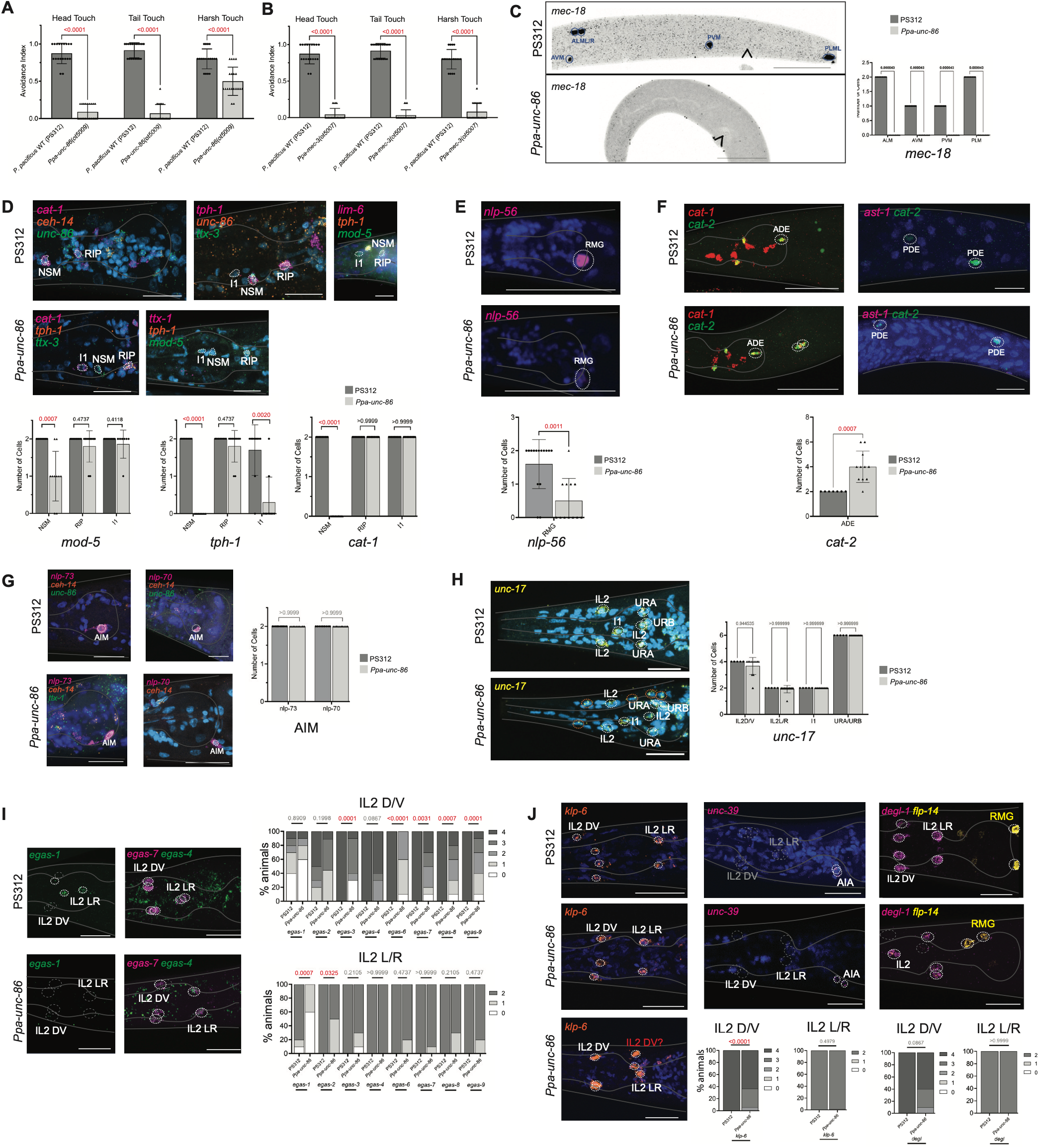
Behavioral and molecular analysis of *Ppa-unc-86* null mutant animals. **(A)** Quantitative analysis of touch assays in *P. pacificus* wild-type (PS312) and *Ppa-unc-86* mutant strains shows touch insensitivity with loss of *unc-86*, as is the case in *C. elegans*. **(B)** Quantitative analysis of touch assays in *P. pacificus* wild-type (PS312) and *Ppa-mec-3* mutant strains shows touch insensitivity with loss of *mec-3*, as is the case in *C. elegans*. **(C – J)** show HCR RNA-FISH staining of various genes as indicated in each panel. **(C)** Loss of *Ppa-unc-86* results in loss of HCR signal of *Ppa-eat-4, Ppa-mec-18*, and *Ppa-mec-3* in the mechanosensory Touch Receptor Neurons (TRNs). Arrow indicates vulva location. **(D)** Loss of *Ppa-unc-86* results in the loss of serotonergic identity of I1 and NSM as indicated by the reduction in *Ppa-tph-1, Ppa-mod-5*, and *Ppa-cat-1*. **(E)** Loss of *Ppa-unc-86* results in the reduction of *Ppa-nlp-56* in RMG. All images shown are single z-slices or projections of >3 z-slices of adult hermaphrodites acquired with confocal microscopy. **(F)** Loss of *Ppa-unc-86* results in ectopic expression of ADE and PDE dopaminergic identity as indicated by additional *cat-1* and *cat-2* HCR signal located posterior of the posterior bulb and in the posterior body wall. These are likely the *unc-86*-expressing neurons AIZ and FLP (near ADE) and PVD (near PDE), which undergo a similar homeotic transformation in *Cel-unc-86* mutants. **(G)** Loss of *Ppa-unc-86* does not result in loss of *Ppa-nlp-73* in AIM. The wild-type panel shown here is the same as shown in Fig.4 and shown for comparison only. **(H)** Loss of *Ppa-unc-86* does not result in loss of *Ppa-unc-17* in *Ppa-unc-86*-expressing neurons of the anterior ganglion (IL2, URA, URB, I1). Ectopic *unc-17* signals are instead observed in presently unidentified cells (circled in orange). **(I)** Loss of *Ppa-unc-86* results in loss of *Ppa-klp-6* and *Ppa-degl* in only IL2 D/V subtype, not IL2 L/R. *Ppa-unc-39*, IL2 subtype selector in *C. elegans*, and *Ppa-flp-14* are not expressed in Ppa IL2 neurons. **(J)** Loss of *Ppa-unc-86* largely results in loss of expression of Ppa *egas* genes across both IL2 subtypes, with a stronger phenotype seen in the IL2 D/V subtype. Scale bars indicate 20 µm. In all quantitative analyses, P-values were calculated by t-test.

Another set of 6-fold radially symmetric ciliated head neurons in the outer labial (OL) sensilla are more distinctly different than the IL neuron subtypes, with different synaptic connectivity, and reflected by distinctive naming in both *C. elegans* and *P. pacificus*: OLQ (“quadrant”) dorsal/ventral neurons and OLL (lateral) neurons (*47*). In *C. elegans*, a Paired homeobox gene, *vab-3/Eyeless*, is expressed in all six OL neurons, whereas a Six-type homeobox gene *ceh-32* is expressed only in the lateral OL (OLL) neurons (*26*). We find the patterns of expression of *P. pacificus vab-3* and *ceh-32* orthologs to be conserved (**Fig. 3H,I**). Other molecular identity features in the subtypes, however, are different: whereas all six OL neurons are glutamatergic in *C. elegans* (*33*), only the OLL neurons are glutamatergic in *P. pacificus* (i.e., express *eat-4/VGluT*; **Fig. 3 H,I**). In contrast, the TRP-type mechanoreceptor channel *ocr-4* is expressed only in OLQ neurons in *C. elegans* (*46, 48*) but its *P. pacificus* orthologue is expressed in all six OL neurons in *P. pacificus* (**Fig. 3 H,I**). Hence, as is the case of the IL2 neurons, the subtype diversification patterns of the labial neurons show evolutionary divergence.

### Amphid sensory neurons reveal many instances of divergence of transcription factor expression

Among the amphid sensory neurons, we observe several differences between *P. pacificus* and *C. elegans*. The ASI and ASG neurons in *P. pacificus*, even though clearly homologs to *C. elegans* ASI and ASG based on anatomy and connectivity (*18, 19*) are each lacking expression of regulatory factors that are critical for their proper development in *C. elegans*. The COE factor *unc-3* is expressed in *C. elegans* ASI where it activates the expression of a number of genes, including the TGFβ homolog *daf-7*, to control dauer formation (*49, 50*). In *P. pacificus*, we fail to observe expression of epitope-tagged *Ppa*-UNC-3 in ASI (**Fig. 2E-H**). Curiously, the *P. pacificus* genome encodes many more *daf-7*-like TGFβ molecules, but none appear to have a role in dauer formation (*51*).

Another amphid sensory neuron class with different expression of a regulatory factor is ASG, which in *P. pacificus* lacks expression of the Pitx-homeobox gene *unc-30* (**Fig. 2I-K**). *C. elegans unc-30* functions in ASG to control innate immunity (*52*), a process that may thereby be regulated in a different manner in *P. pacificus*.

The terminal selector of the *C. elegans* ASE gustatory neuron pair, the zinc finger transcription factor *che-1* (*53*), is also expressed in the *P. pacificus* ASE neurons (**Fig. 3J**) (*19, 54*). Glutamatergic identity of the ASE neurons (expression of *eat-4/VGluT)* is also preserved in *P. pacificus* (**Fig. 3J**). In *C. elegans*, these neurons are laterally diversified by asymmetric expression and function of the homeobox gene *lim-6* in ASEL (*55, 56*). Even though the ASE neurons also appear to be functionally lateralized in *P. pacificus* (*19, 47*), we noted that *Ppa*-*lim-6* is expressed in both neurons of this pair, indicating that the lateralization of the Ppa-ASE neurons must be modulated by a different regulator (**Fig. 3J**).

Also, as previously reported (*57*), we observed a “gain” of *che-1* expression in the AFD thermosensory neuron of *P. pacificus* (note that we use the term “gain” here, and in the ensuing sections, just in comparison to *C. elegans;* in an evolutionary sense, it is equally possible that the expression of *che-1* in AFD is ancestral and was lost in *C. elegans*). We assessed to what extent the *P. pacificus* AFD neurons retain co-expression of two known *C. elegans* AFD identity regulators, the Otx-type homeobox gene *ttx-1* and the LIM homeobox gene *ceh-14* (*58, 59*). We find that *Ppa-ttx-1* and *Ppa-ceh-14* indeed show conserved expression in Ppa-AFD (**Fig. 3K**). Ppa-AFD also displays a glutamatergic and cholinergic co-transmitter identity (**Fig. 3K,L**), like its *C. elegans* counterpart.

We find that the Prop-type homeobox gene *unc-42* is expressed in ASH neurons in both *C. elegans* and *P. pacificus*, and this neuron pair is also glutamatergic in both species (**Fig. 2L,M, Fig. 8C**). However, *Ppa-unc-42* is also expressed in an additional as yet unidentified, *P. pacificus* sensory neuron bilateral pair, as assessed by (a) cell counts of epitope-tagged *Ppa*-UNC-42 (**Fig. 2L,M**) and (b) co-expression of the pan-sensory maker *osm-6* with *unc-42* by HCR in two pairs of cells in *P. pacificus* instead of a single sensory neuron pair (ASH) in *C. elegans* (**Fig. 8C**).

**Figure 8.**
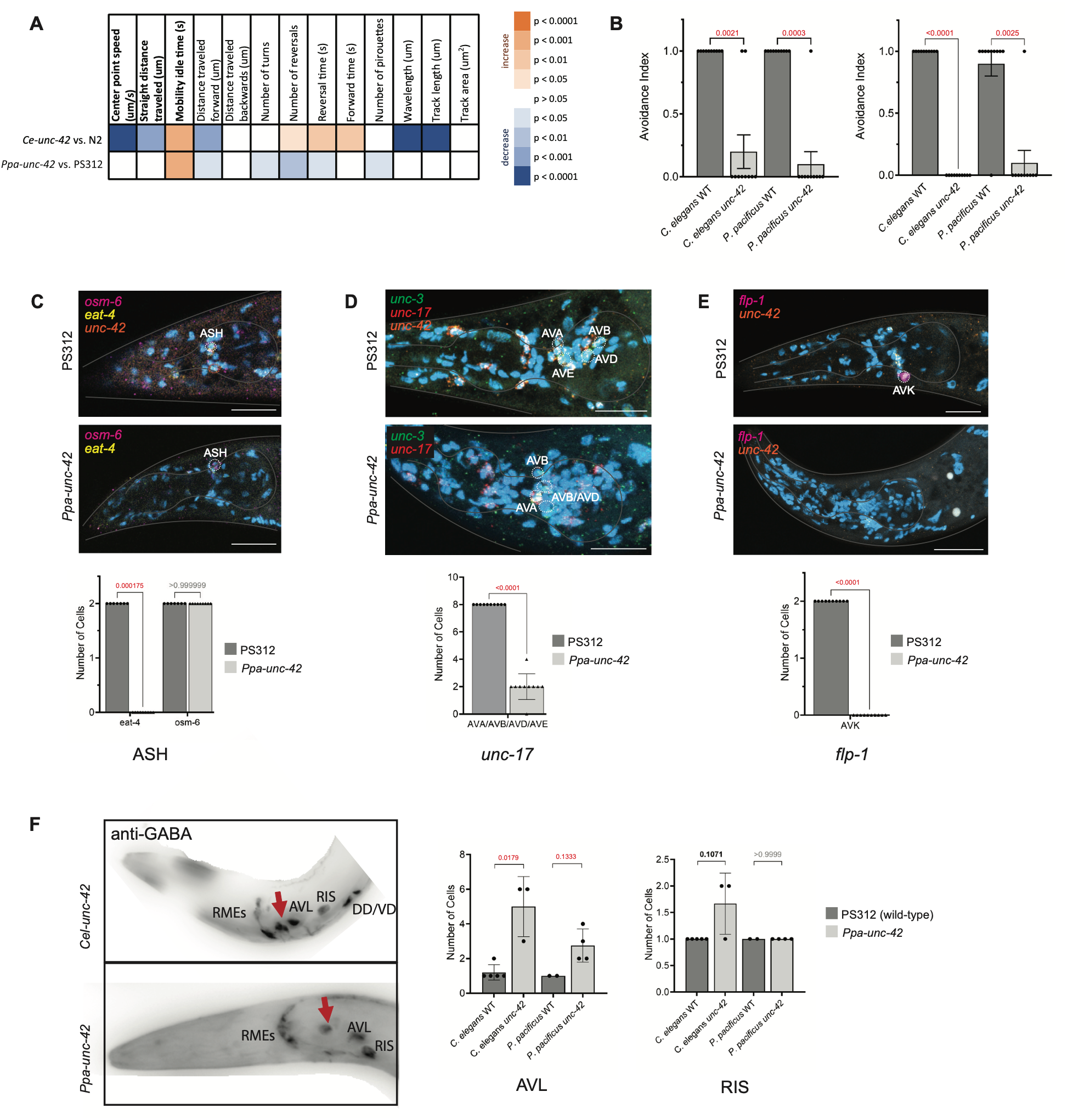
Behavioral and molecular analysis of *Ppa-unc-42* null mutant animals. **(A)** Heatmap showing comparison of locomotory characteristics between wild-type *C. elegans* (N2), *Cel-unc-42* null mutant, wild-type *P. pacificus* (PS312) and *Ppa-unc-42* null mutant. Orange indicates positive significant change, and blue indicates negative significant change. Loss of *unc-42* in both species results in a reduction of locomotion and increase in mobility idle time as expected of the ‘unc’ phenotype. **(B)** Quantitative analysis of nociceptive touch and octanol assays show that loss of *unc-42* results in loss of nociception in both species. **(C)** Loss of *Ppa-unc-42* leads to loss of terminal identity marker *Ppa-eat-4* in ASH **(E)** Loss of *Ppa-unc-42* leads to loss of *Ppa-unc-17* in AVA/AVB/AVD/AVE **(E)** Loss of *Ppa-unc-42* leads to loss of *Ppa-flp-1* in AVK. **(F)** Loss of *unc-42* results in ectopic anti-GABA in head neurons in *P. pacificus* and *C. elegans*. Ectopic cells are more commonly seen near AVL than RIS in *P. pacificus*. All images shown are single z-slices or projections of >3 z-slices of adult hermaphrodites acquired with confocal microscopy. Scale bars indicate 20 µm. All P-values calculated by t-test or one-way ANOVA.

The glutamatergic *C. elegans* ASK sensory neuron class is specified by the *mls-2* Hmx-type homeobox gene and the *ttx-3* LIM homeobox gene (*25, 33*). As mentioned above, in *P. pacificus*, the *mls-2* gene has triplicated. One copy, *mls-2.1* shows no expression anywhere, but both *mls-2.2* and *mls-2.3* are co-expressed with the *eat-4/VGluT* marker in the ASK neurons (**Fig. 3M,N**). However, while its expression in interneurons is conserved (discussed further below), *Ppa-ttx-3* fails to be expressed in ASK.

We also examined the bilateral AUA neurons, which are not amphid neurons, but send a dendrite along the amphid dendrites towards the nose, albeit not completely, in both *C. elegans* and *P. pacificus* (*19, 47*). In other more basal nematodes, this neuron appears to be a functional amphid neuron with the dendrite extending to the tip of the nose (*60*). *C. elegans* AUA is glutamatergic and requires the paralogous, adjacent Olig-like bHLH genes *hlh-17* and *hlh-32* for proper differentiation (*61*). In *P. pacificus, hlh-17* has not duplicated and we find that *P. pacificus* AUA co-expresses *Ppa eat-4/VGLUT* and *Ppa-hlh-17* (**Fig. 4A**), indicating that the morphological similarity between *Cel* and *Ppa* AUA extends to key molecular features.

### Conservation and divergence among head interneurons

Moving beyond its expression in sensory neurons, *unc-86* expression is also observed in a diverse set of interneurons in *C. elegans*, including the AIZ, AIM, RIH and RIP neurons (*30*). Antibody staining, as well as HCR, reveals *Ppa-unc-86* expression in cells of similar location in *P. pacificus* (**Fig. 2A,B, Fig. 4**). The *P. pacificus* neuron that is in a position similar to AIZ also expresses the Dlx ortholog *ceh-43*, previously shown to cooperate with UNC-86 in controlling AIZ identity in *C. elegans* (**Fig. 4B**)(*26*). Moreover, *Ppa-eat-4/VGluT* expression indicates that AIZ is also glutamatergic, as it is in *C. elegans*. The ventral midline unpaired RIH interneuron is also UNC-86-positive in both species and *P. pacificus* RIH expresses, like in *C. elegans*, a conserved neuropeptide, *nlp-71* (**Fig. 4C**).

The RIP interneuron, the sole connector of the central and enteric nervous systems in both *C. elegans* and *P. pacificus* (*18*), is serotonergic in *P. pacificus* (*37*) but not *C. elegans*, an observation we confirm here by HCR (**Fig. 4D)**. Also in contrast to *C. elegans*, we detect expression of GABA synthesis and vesicular transport genes (*unc-25/GAD* and *unc-47/VGAT*) in *Ppa*-RIP (**Fig. 4D**). In *C. elegans*, RIP is cholinergic, and we find in *P. pacificus* that along with expression of a serotonergic identity, we observe lack of the cholinergic marker *unc-17/VAChT* expression by HCR, and, similarly, no expression of a conserved neuropeptide-encoding gene, *nlp-73* expressed in *C. elegans* RIP (**Fig. 4E,F;** in contrast, *nlp-73* expression in AIM is conserved between both species). Despite these notable changes in neuronal features, we find the presently known regulatory signature of RIP to be conserved as Ppa-RIP still co-expresses *unc-86* and the *ttx-1* Otx-type homeobox genes (**Fig. 4E,G**), which control RIP differentiation in *C. elegans* (*26*). Whether other regulatory factors that may cooperate with *unc-86* and *ttx-1* exist and have diverged between these two nematodes, remains an open question.

The bilateral AIM interneurons, which are glutamatergic in *C. elegans*, also shows patterns of divergence. The Ppa-AIM interneurons does not show the *eat-4/VGLUT* expression we observed in *C. elegans* (**Fig. 4H-J**), nor does it show serotonin staining or *mod-5/SERT*-dependent serotonin uptake as observed in *C. elegans* (*37*). Yet we find that the *P. pacificus* orthologs of two conserved neuropeptides, *nlp-70* and *nlp-73* that are prominently expressed in *C. elegans* AIM (*26, 46*), are still expressed in the AIM neurons of *P. pacificus* (**Fig. 4J,E**). Also, in striking similarity to *C. elegans* (*62*), the AIM neuron still adopts a cholinergic phenotype, i.e., *unc-17/VAChT* expression, exclusively in male animals (**Fig. 4K**).

Two terminal selectors of AIM fate in *C. elegans, unc-86* and the LIM homeobox gene *ceh-14* (*33*), are also expressed in AIM in *P. pacificus* (**Fig. 4H-K**). A third factor in this regulatory signature, the Hmx-type homeobox gene *mls-2* (*26*) triplicated in *P. pacificus* to generate three *mls-2* loci (**Fig. 1**). While the expression of two of them is conserved in the ASK and AVJ neurons (**Fig. 3M**), neither shows robust expression in AIM, indicating shift in the regulatory signature of AIM.

Two amphid interneuron classes, AIY and AIA, are cholinergic in *C. elegans* and specified by the LIM homeobox gene *Cel-ttx-3*, cooperating with the Prd-type *Cel-ceh-10* homeobox gene in AIY and the Six-type homeobox gene *Cel-unc-39* in AIA (*26, 63*). We find both neuron classes to also be cholinergic in *P. pacificus* and to retain the regulatory signatures of the *P. pacificus* orthologs *ttx-3* with *ceh-10* in AIY and *ttx-3* with *unc-39* in AIA (**Fig. 4L,M**).

Another set of interneurons, anatomically defined in both *C. elegans* and *P. pacificus*, are the ‘command interneurons’ (AVA, AVB, AVD, AVE) that relay various sensory inputs to generate appropriate motor outputs via ventral nerve cord motor neurons. The cholinergic identity of these neurons is evident in *P. pacificus* as well (**Fig. 8D**). In *C. elegans*, the unique combinatorial signature that functionally defines these command interneurons is the Prop1-type homeobox gene *unc-42* and the COE-type transcription factor *unc-3* (*64, 65*). Both antibody staining and HCR co-staining show that *P. pacificus unc-42* and *unc-3* are co-expressed in the *P. pacificus* command interneurons as well (**Fig.2L,M, Fig. 4N, Supp. Fig. S3**). Other cholinergic interneurons in the head include sublateral interneurons SIB and SAA, which require Cel-UNC-42 for their proper differentiation (*65*). Ppa-UNC-42 antibody staining shows expression in cells that by position can also be identified is the SIB and SAA neurons **(Fig. 2L,M**).

Lastly, we examined the single octopaminergic interneuron pair found in *Cel* and *Ppa*, RIC, identified by the unique co-expression of synthetic enzyme genes *tdc-1* and *tbh-1* (*37, 66*). In *C. elegans*, RIC identity is defined by the deeply conserved bHLH transcription factor HLH-13, the ortholog of the vertebrate PTF1a and NATO3 genes (*61*). *C. elegans hlh-13* is exclusively expressed in RIC (*61*); we found that the sole *Ppa-hlh-13* ortholog is also expressed only in the RIC interneuron in *P. pacificus* (**Fig. 4O**).

### Neuropeptidergic interneurons show conserved and novel signatures

Three interneuron classes in the *C. elegans* head, ALA, AVK and RMG, do not appear to use classic small molecule neurotransmitter, as assessed by lack of expression of vesicular transporters and synthesizing enzymes for acetylcholine, GABA, glutamate or monoamines (*67*). Instead, all three different neuron classes express a multitude of distinct neuropeptides (*46*) to control various homeostatic processes. For each of the three neuron classes, specific combinations of transcription factors have been found to control their identity in *C. elegans: ceh-14* in ALA (*68*), *unc-42* in AVK (*69*) and *unc-86* in RMG (*26*).

The peptidergic identity of *C. elegans* AVK is manifested by co-expression of many neuropeptides, including the AVK-exclusive expression of *Cel-flp-1* (*46, 69*). In *P. pacificus*, AVK is also the only neuron expressing the *Ppa-flp-1* ortholog (**Fig. 8E**); and like in *C. elegans* it lacks expression of *unc-17, eat-4, unc-47* and monoaminergic synthesis machinery, hence displaying the same “neuropeptidergic-only” phenotype as in *C. elegans*. Also like in *C. elegans*, the *Ppa-unc-42* homeobox gene, the terminal selector of AVK identity (*69*), is expressed in *P. pacificus* AVK (**Fig. 2L,M, Fig. 8E**).

The peptidergic ALA neuron, which controls sleep behavior in *C. elegans*, expresses the sleep-controlling *flp-13* (*70*) and its identity is controlled by the LIM homeobox gene *ceh-14* (*68*). We found that the *P. pacificus* ALA neuron, like in *C. elegans*, a large unilateral neuron with a distinctive position dorsal to the nerve ring, also co-expresses *Ppa-ceh-14* and *Ppa-flp-13* (**Fig. 4P**)

*P. pacificus* RMG neurons also expresses the homeobox gene *unc-86* (**Fig. 2A,B**), as well as *P. pacificus* orthologs of the neuropeptides *flp-14* and *nlp-56* that are prominently expressed in *C. elegans* RMG (**Fig. 4Q**)(*26, 46*). Intriguingly, *Ppa-*RMG appears to adopt a glutamatergic identity, and appears to do so in a lateralized manner: it is consistently the right RMG neuron (RMGR), but not RMGL that expresses *eat-4/VGluT*, the indicator of glutamate usage (**Fig. 4Q**).

### Interneurons in the retrovesicular ganglion and tail show conserved molecular signatures

The retrovesicular ganglion demarcates the anterior end of the ventral nerve cord. Apart from harboring several ventral nerve cord motor neurons that we will discuss in the ensuing sections, the *C. elegans* retrovesicular ganglion contains two interneuron classes that we examined for phylogenetic conservation. One is the RIG interneuron pair, the only glutamatergic neurons in the retrovesicular ganglion of *C. elegans* (*33*), which also expresses the LIM homeobox gene *Cel-lim-6* (*55*). We find that, similarly, in *P. pacificus* a single glutamatergic (i.e. *eat-4/VGLUT-*positive) neuron pair is present in the retrovesicular ganglion, and it expresses *Ppa-lim-6* (**Fig. 4R**), indicating conservation of the key identity features of this neuron class.

Another interneuron class in the *C. elegans* retrovesicular ganglion is the AVF pair, which is thought to be involved in communicating with the mid-body HSN neuron to modulate egg-laying behavior (*71*). This neuron class does not synthesize its own conventional neurotransmitter in *C. elegans*, but serves as an apparent sink of GABA, taking up this neurotransmitter via the GABA uptake transporter SNF-11 (*72*). This function is controlled by the AVF-expressed Prd-type homeobox gene *unc-4* in *C. elegans* (*72*). We find that co-expression of *Ppa-unc-4* and *Ppa-snf-11* is preserved in the retrovesicular ganglion in *P. pacificus* (**Fig. 4S**).

### Midbody neurons are largely conserved

Apart from the above-mentioned UNC-86/MEC-3-dependent touch receptor neurons and dopaminergic AST-1/CEH-43-positive PDE mechanosensory neurons, *C. elegans* also contains other midbody neurons, including the Q neuroblast-derived SDQ neurons, the BDU interneuron pair, the CAN neurons and the HSN neurons (*47*).

The *Ppa* CAN neurons were previously visualized in *P. pacificus* via the expression of a *P. pacificus-*specific gene, dauerless (*73*). The role of CAN as the central production hub of a class of internal signaling molecules, the nemamides (*74*), appears to be conserved. A ortholog of an enzyme that catalyzes their production in *C. elegans* (e.g. the non-ribosomal peptide synthetase *nrps-1)*(*74*) is present in the *P. pacificus* genome and transcripts for *Ppa-nrps-1* are exclusively found in Ppa-CAN (**Fig. 5A**). The regulatory signature of CAN also appears to be preserved. In *C. elegans*, this regulatory signature is defined by the unique combination of two homeobox genes, the Prd-type homeobox gene *Cel-ceh-10* and the Distalless/Dlx ortholog *Cel-ceh-43*, both of which are required for CAN neuron differentiation (*75–77*). We find that the Ppa-CAN neuron expresses the same combination of homeobox genes (**Fig. 5B**).

The existence and position of BDU, SDQ and HSN homologs in *P. pacificus* remained unresolved, since our previous serial section EM reconstruction only covered the head region of the worm. Lack of serotonin-staining in the midbody of *P. pacificus* also questioned the existence of the HSN neurons, which exists and is serotonergic in a range of nematode species (*78*).

A panneuronal probe (*Ppa-egl-3*) labels a set of midbody neurons including the touch neurons, as well as neurons whos position is consistent with being the *P. pacificus* BDU and SDQ neurons (**Fig. 5C**). Using UNC-86 antibody staining (**Fig. 2A-D**), as well as HCR probes against other transcription factors previously described to be expressed in these midbody neurons in *C. elegans*, supports these neurons as being the *P. pacificus* homologs of the BDU and SDQ neurons. Specifically, a neuron pair in the anterior half of the animal co-expresses the *P. pacificus* orthologs of the regulatory signature characteristic of the *C. elegans* BDU neuron (*77*), namely *Ppa-unc-86*, the LIM homeobox gene *Ppa-ceh-14* and the Dlx ortholog *Ppa-ceh-43* (**Fig. 5D,E**). While *C. elegans* expresses no BarH/Barx-like CEH-30 protein in BDU (*77*), *Cel-ceh-30* transcripts are detected in BDU in scRNA datasets and we detect transcripts for one of the two *P. pacificus* BarH/Barx-like homeobox gene in BDU as well (*ceh-31.2)*(**Fig. 5D,E**). While *C. elegans* BDU neurons are peptidergic and do not employ a small, classic neurotransmitter, *P. pacificus* BDU appear to be cholinergic, as inferred from *Ppa-unc-17/VAChT* expression (**Fig. 5C**).

The SDQL/R neuron pair, lineal sisters of UNC-86-expressing AVM and PVM mechanosensory neurons (*79*), are a pair of two interneurons of presently unknown function. In *C. elegans* these neurons are (a) cholinergic (*unc-17/VAChT-*positive) and (b) defined by the combinatorial expression of *ceh-43/Dll* and *ceh-31/Barx* homeobox genes (*77*). We find *P. pacificus* neurons with a left/right asymmetric position similar the Cel-SDQ neurons (SDQR anterior to the vulva and SDQL posterior to the vulva), displaying the same neurotransmitter phenotype and *ceh-43 & ceh-31-*regulatory signature (**Fig. 5C,F**). Both *P. pacificus* duplicates of an apparently ancestrally single *ceh-31* gene, *Ppa-ceh-31.1* and *Ppa-ceh-31.2*, are co-expressed in SDQ.

Another bilateral midbody neuron pair, the hermaphrodite-specific HSN, is defined in *C. elegans* by the combinatorial expression of a number of transcription factors, including UNC-86, AST-1 and the bHLH protein HLH-3 (*80, 81*). In *C. elegans*, HSN deploys two neurotransmitter, serotonin and acetylcholine (*67*). No serotonergic neuron is observed in the midbody of *P. pacificus* (*78*), but we detected a cholinergic (i.e. *unc-17*-positive) neuron pair in *P. pacificus* in a position slightly more posterior to where *C. elegans* HSN is located (**Fig. 5G,H**). This neuron pair coexpresses *Ppa-unc-86* (strongly) and *Ppa-ast-1* (weakly) but lacks expression of *Ppa-hlh-3* (**Fig. 5G,H**)(since our *Ppa-hlh-3* HCR probes detect no signal anywhere, the absence in HSN may be caused by technical reasons). Consistent with this UNC-86-positive neuron being the *P. pacific* homolog of HSN, we do not observe UNC-86 antibody staining in the midbody section of males (**Supp.Fig. S4**), in which, in *C. elegans*, HSN is removed by sex-specific apoptotic cell death (*82*).

### Conservation of cholinergic ventral nerve cord motor neuron classes

Within the ventral nerve cord, we find through anti-GABA antibody staining, as well as *unc-17/VAChT* and *unc-25/GAD* expression that the *P. pacificus* ventral nerve cord is populated, like in *C. elegans*, by distinct GABAergic and cholinergic motor neurons (**Fig. 6B**). In *C. elegans* the identity of cholinergic motor neurons is controlled by the COE-type transcription factor *unc-3* (*50, 83*). In *P. pacificus*, Ppa-UNC-3::2xFLAG antibody staining reveals many ventral nerve cord motor neurons (**Fig. 2F-H**). Their overall count and patterns is similar to the Cel-UNC-3 expression in cholinergic ventral nerve cord motor neurons. Co-HCR against *Ppa-unc-17/VAChT* and *Ppa-unc-3* in adults shows that all *unc-3*-positive neurons in the ventral nerve cord are cholinergic (shown in **Fig. 9** in the context of *unc-3* mutant analysis).

There are several distinct classes of cholinergic, UNC-3(+) ventral nerve cord motor neurons in *C. elegans*, including dorsal and ventral A types (DA and VA), dorsal and ventral B types (DB, VB) and AS types. Each of these can be identified via class-specific transcription factor codes that cooperate with UNC-3 to shape their unique identities (*84*): DA/VA-types and DB/VB-types are distinguished by expression of the DA/VA-specific *unc-4* homeobox gene (*85*) and the DB/VB-specific *ceh-12* homeobox gene (*86*), VA/VB are distinguished from DA/DB by the VA/VB-specific *bnc-1* Zinc finger transcription factor (*84*), DB from VB by the DB-specific Eve-type *vab-7* homeobox genes (*87*). The *C. elegans* AS neurons uniquely co-express the orphan receptor *unc-55* and the T-box gene *mab-9*, which is also expressed in DA and DB (*88, 89*). Based on combinatorial co-staining with specific HCR probe sets, we detect similar expression patterns of their *P. pacificus* orthologs: *Ppa-unc-4* is expressed in DA/VA, *Ppa-ceh-12* in VB, *Ppa-vab-7* in DB, *Ppa-bnc-1* in VA and VB, and *Ppa-unc-55* and *Ppa-mab-9* in AS (and DA and DB)(**Fig. 9E-H**).

**Figure 9.**
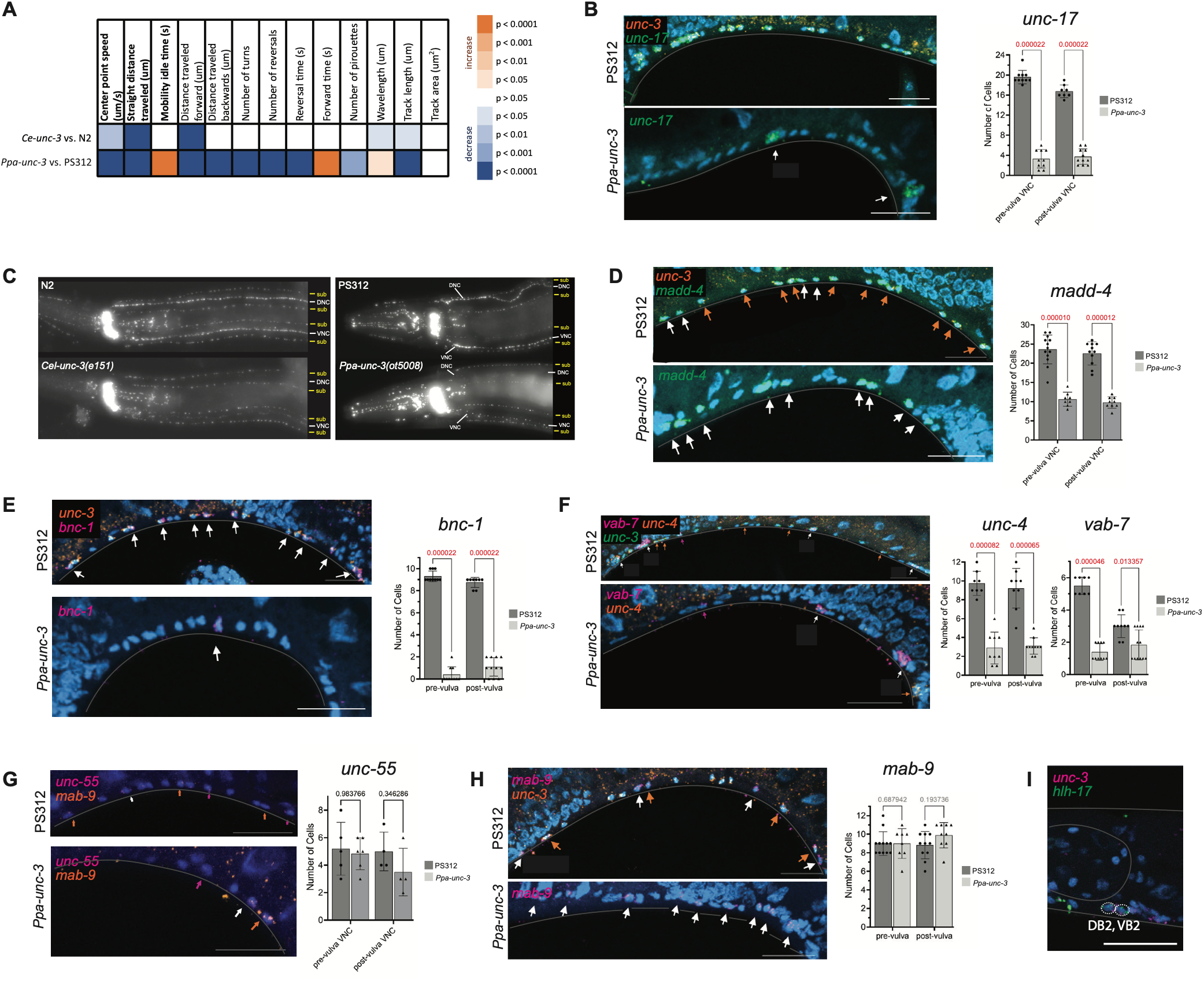
Behavioral and molecular analysis of *Ppa-unc-3* null mutant animals. **(A)** Heatmap showing comparison of locomotory characteristics between wild-type *C. elegans* (N2), *Cel-unc-3* null mutant, wild-type *P. pacificus* (PS312) and *Ppa-unc-3* null mutant. Orange indicates positive significant change, and blue indicates negative significant change. Loss of *unc-3* in both species results in a reduction of locomotion and increase in mobility idle time as expected of a severe ‘unc’ phenotype. **(B – H)** show HCR RNA-FISH staining of various genes as indicated in each panel. **(B)** *Ppa-unc-3* regulates the cholinergic terminal identity of neurons in the VNC as indicated by the loss of expression of *Ppa-unc-17* in the absence of *Ppa-unc-3*. Arrows indicate the VC neurons, which express *Ppa-unc-17* but do not express *Ppa-unc-3* and are not affected by the null mutation. **(C)** Loss of *Ppa-unc-3* results in the reduction of *Ppa-madd-4* in the VNC neurons. Orange arrows indicate *unc-3*/*madd-4* co-expressing neurons, and white arrows indicate neurons that only express *madd-4*. **(D)** Loss of *Ppa-unc-3* results in the reduction of *Ppa-unc-4* and *Ppa-vab-7*, which are expressed in the VA/DA (orange arrows) and DB (magenta arrows) neurons, respectively, and are co-expressed (orange arrows) in the VC neurons of the VNC. **(E)** Loss of *Ppa-unc-3* results in the reduction of *Ppa-bnc-3* in the VA/VB neurons (white arrows) in the VNC. **(F)** Loss of *Ppa-unc-3* does not result in the loss or reduction of *Ppa-mab-9* in the AS/DA/DB neurons of the VNC. (Orange arrows indicate *unc-3/mab-9* co-expressing neurons, presumably the AS/DA/DB neurons, and white arrows indicate neurons only expressing mab-9.) **(G)** Loss of *Ppa-unc-3* does not result in the reduction of *Ppa-unc-55* in the AS neurons in the VNC. **(H)** Loss of *Ppa-unc-3* results in the reduction of *Ppa-eat-4* in DVC. **(I)** Loss of *unc-3* in both *C. elegans* and *P. pacificus* results in the reduction of anti-ACh signal in the ventral and dorsal nerve cords, consistent with the loss of cholinergic identity of the neurons in the VNC.

In *C. elegans*, individual members of the A- and B-type motor neuron classes are further diversified into individual subtypes (*90, 91*). For example, the DB2 and VB2 B-type member class members, located in the retrovesicular ganglion are distinguishable from other, ventral cord-localized B-type members by the expression of the paralogous and genomically adjacent, Olig-like *hlh-17* and *hlh-32 C. elegans* bHLH genes (*61*). In *P. pacificus hlh-17* has not duplicated and its sole representation is also expressed in two, *unc-3/EBF* and *unc-17/VAChT-*positive neurons of the retrovesicular ganglion, akin to the DB2 and VB2 in *C. elegans* (**Fig. 9I**).

The only other ventral nerve cord cholinergic neuron class that does not express *C. elegans* or *P. pacificus* UNC-3 are the VC neurons. In contrast to *C. elegans* in which only a subset of VC neurons sporadically take up serotonin via the *mod-5* transporter (*92*), all four *P. pacificus* VC neurons synthesize their own serotonin (*37, 78*). We find that as in *C. elegans*, the *Ppa-*VC neurons co-express the *unc-4* and *vab-7* homeobox genes (**Fig. 5I**; VC neurons are identified by co-staining with *cat-1/VMAT*), but unlike *C*. elegans, they do not express the bHLH gene *hlh-3*, which has been implicated in VC identity specification in *C. elegans* (*93*). Hence, among the ventral cord motor neurons, based on the markers we analyzed, the VC neurons appear to display the most diversification of fate markers (one transcription factor and neurotransmitter identity).

Cholinergic motor neurons are not only present in the ventral nerve cord, but also in the head ganglia where they innervate head muscle. In *C. elegans*, most of these require the *Cel-unc-42/Prop1* homeobox gene for their proper differentiation (RMD, SMD, RMF, RMH) (*65*). Through antibody staining we observe expression of Ppa-UNC-42 in these neuron classes as well (**Fig. 2L,M**).

### Conservation and novelties of GABAergic ventral nerve cord motor neuron classes

Like in *C. elegans*, D-type motor neurons of *P. pacificus* stain with anti-GABA antibody (**Fig. 6A**) and they express key components of the GABA synthesis and transport pathway, *unc-25/GAD, unc-47/VGAT* and *unc-46/LAMP* (**Fig. 6, 12**)(*94*). They interdigitate, like in *C. elegans* with cholinergic motor neurons of the ventral nerve cord (**Fig. 6B**). Expression of other molecular identity features of *C. elegans* D-type motor neurons is preserved in *P. pacificus*, including genes encoding the synaptic organizer, *madd-4*, the phosphodiesterase *pde-4* or the lysozyme *ilys-4* **(Fig. 11F**).However, the *P. pacificus* homolog of a *C. elegans*, D type-motor neuron-expressed nicotinic acetylcholine receptor, *acr-14*, that possibly mediates input from cholinergic command interneurons (*95*), is not expressed in the *P. pacificus* D-type neurons.

**Figure 10.**
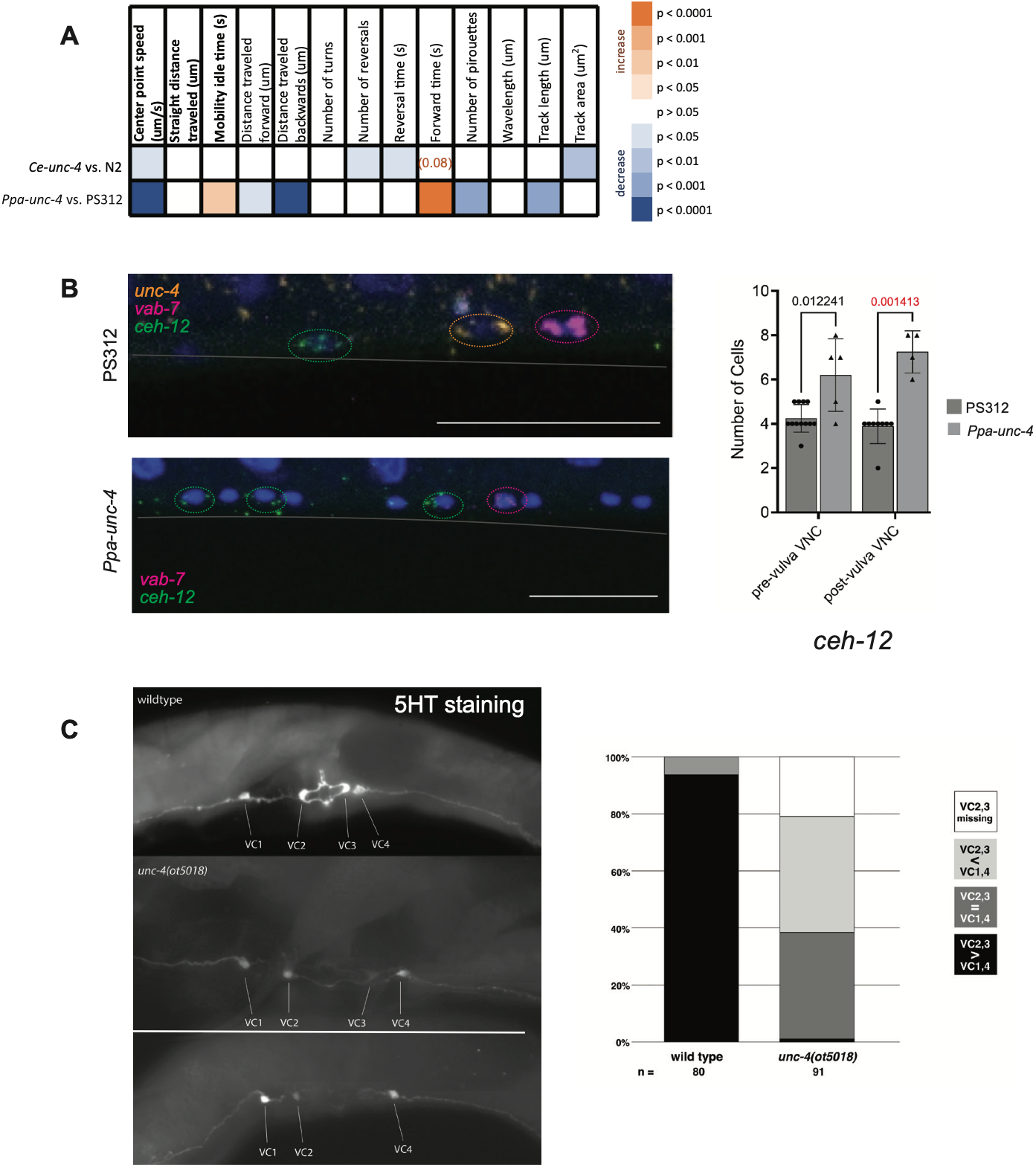
: Behavioral and molecular analysis of *Ppa-4 unc-4* mutant animals. **(A)** Heatmap showing comparison of locomotory characteristics between wild-type *C. elegans* (N2), *Cel-unc-4* null mutant, wild-type *P. pacificus* (PS312) and *Ppa-unc-4* null mutant. Orange indicates positive significant change, and blue indicates negative significant change. Loss of *unc-4* in both species results in a reduction of locomotion as expected of a ‘unc’ phenotype. **(B)** Loss of *Ppa-unc-4* results in the derepression of *Ppa-ceh-12* in DA/VA neurons of the VNC. **(C)** Serotonin-antibody staining of *unc-4* mutants. Scale bars indicate 20 µm. All P-values calculated by t-test.

The identity of GABAergic D-type neuron in the ventral nerve cord of *C. elegans* is controlled by the Pitx-like homeodomain protein UNC-30 (*95–97*). Antibody staining of epitope-tagged Ppa-UNC-30, as well as *unc-30* HCR, combined with either GABA antibody staining or HCR against key components of the GABA synthesis and transport pathway (*unc-25/GAD* and *unc-47/VGAT)*, shows that the expression of this *C. elegans* terminal selector is conserved in *P. pacificus* (**Fig. 2I-K, Fig. 11D**). However, a co-factor of *C. elegans unc-30*, the GATA transcription factor *elt-1* (*72*), has duplicated in *P. pacificus*, but neither of the two genomically adjacent *Ppa-elt-1* paralogs shows expression in GABAergic ventral nerve cord neurons (**Fig. 11I**). Instead, *Ppa-elt-1.2* shows expression in a subset of *unc-3*-positive ventral nerve cord neurons (**Fig. 11I**).

The GABAergic D-type neurons come in two distinct, dorsally projecting (DD) and ventrally projecting (VD) subtypes. The differences between these subtypes are controlled by the orphan nuclear receptor UNC-55 in *C. elegans*, which is expressed in VD neurons to repress DD features (*88*). We find that the *Ppa-unc-55* gene is also expressed in only a subset of D-type neurons, which we assume to be the VD neurons (**Fig. 11H**). The type of molecular features that distinguish DD and VD have, however, diversified, as exemplified by two genes that are expressed in a subtype-specific manner in *C. elegans*. HCR probes against the *P. pacificus* ortholog of the *flp-13* gene, a DD subtype-specific marker in *C. elegans*, show no signals in the Ppa-D-type neurons. The *oig-1* gene, a synaptic organizer restricted to the VD-type neurons in adult animals (*98*), is genomically absent in *P. pacificus*.

### GABAergic inter- and motor neurons show conserved regulatory signatures

Apart from the GABAergic motor neurons in the ventral nerve cord, there are five other classes of GABA synthesizing and secreting neurons in *C. elegans*. Three of them are motor neurons, one of which is a head motor neuron class (RME), the other two are hindgut musculature-innervating motor neurons (AVL, DVB). The fourth and fifth class are head interneurons RIS and RIB (*99*). Anti-GABA staining and HCR against *unc-25/GAD* transcripts reveals that these neurons are also GABAergic in *P. pacificus* (**Fig. 6C; Fig. 11D**)(*94*). We further corroborated this notion through the generation of transgenic *P. pacificus* strains that fused parts of the *unc-25/GAD* locus to RFP (**Fig.6D**).

In *C. elegans*, the LIM homeobox gene *lim-6* acts as a terminal selector of the RIS, AVL and DVB GABAergic neurons and is also expressed in a subset of the RME motor neurons, the RME lateral (L/R) pair (*55, 72*). HCR probes against *lim-6*, combined with *unc-25/GAD* probes show that *Ppa-lim-6* is expressed in the same set of neurons (**Fig. 6C**). In the RIB neurons, *Cel-unc-25* expression overlaps with *Ce-ttx-1* homeobox gene expression, and this overlap is preserved in *P. pacificus* (**Fig. 4L**). Taken together, the identity and regulatory signatures of head, ventral cord and tail GABAergic neurons appear to be conserved.

### Conservation and novelties in tail neurons

In *C. elegans*, the PVP interneuron pair, located in the pre-anal ganglion, shows a unique overlap in the expression of *unc-3* and *unc-30* and they cooperate to control the cholinergic identity of PVP (*64*). We observe a similar molecular signature in a preanal neuron pair, which we assume to be the Ppa-PVP neurons: This pair is cholinergic (i.e. *unc-17* positive) and co-expresses *Ppa-unc-3* and *Ppa-unc-30* (**Fig. 11E**).

Further back in the tail, a unilateral, glutamatergic interneuron, PVR, which also sends a process into the nerve ring in both *P. pacificus* and *C. elegans* (*18*), shows conserved molecular features. Like in *C. elegans*, it expresses both *Ppa-eat-4/VGluT* and both copies of the *Ppa-*BarH1 homologs, *ceh-31.1* and *ceh-31.2* (**Fig. 4T,U**). The co-expression of *ceh-31.1* and *ceh-31.2* invites a fascinating comparison of the expression of these two *P. pacificus* duplicates compared to the independently duplicated *ceh-31* and *ceh-30* genes (summarized in **Table S2**): The expression of all 4 genes is preserved in SDQ. Compared to *C. elegans ceh-31* (the ancestral version of the gene, based on sequence analysis; **Fig. 1**), the two *P. pacificus* homologs also preserve expression in PVR, yet only one of them (*ceh-31.2)* preserves expression in URB and BDU, while apparently losing expression in URA and AVB.

### Conserved regulatory signatures in pharyngeal enteric neurons

The pharyngeal enteric nervous system of both nematode species is composed of a homologous set of 14 neuron classes (*100*). One neuron class whose identity regulation is particularly well understood in *C. elegans*, is the serotonergic NSM neuron, in which two homeobox genes, *unc-86* and *ttx-3* cooperatively define serotonergic NSM identity (*41, 101*). Anti-serotonin staining in *P. pacificus* also labels the NSM neurons and these cells express serotonergic marker genes such as *tph-1* and *mod-5* (*37, 78*). We found that *P. pacificus* also shows a unique overlap of *Ppa-unc-86* and *Ppa-ttx-3* in the NSM neuron (**Fig. 4V**). Like in *C. elegans*, anti-UNC-86 also shows staining in *P. pacificus* I1 (**Fig. 2A,B**), which co-expresses *tph-1* (**Fig. 7D**).

### Male-specific CEM and MCM neurons appear to be conserved

The nervous system of male nematodes contains additional neurons that are involved in various aspects of mate sensing and copulatory behavior. *C. elegans* generates male-specific neurons through three distinct processes, sex-specific blast cell proliferation, sex-specific cell death and sex-specific transdifferentiation (*79, 102*). To what extent these processes are conserved in *P. pacificus* is not known. We examined two specific paradigms, the male-specific transdifferentiation of a glial cell that generates the male-specific neuron, MCM (*102*) and a hermaphrodite-specific cell death that results in the male-specificity of the CEM neurons (*79*). For both neuron classes, terminal selectors have been identified, *unc-86* for the CEM neurons, and *unc-42* for the MCM neurons. We observed male-specific *Ppa-unc-86* and *Ppa-unc-42* signals in the head of male animals, with their position and number (4 CEM and 2 MCM) being consistent with the presence of Ppa-CEM (UNC-86-expressing) and Ppa-MCM neurons (UNC-42 expressing) (**Supp. Fig. S4**).

### Conservation of panneuronal molecular regulator signature

In *C. elegans*, extensive expression pattern analysis has shown that all neurons share several “pan-neuronal” molecular features, including genes involved in the synaptic vesicle cycle and genes involved in neuropeptide processing and transport (*103*). We generated HCR probes against one member of each functional category, *Ppa-rab-3* (encoding a synaptic vesicle protein) and *Ppa-egl-3* (neuropeptide processing). Both show pan-neuronal expression (**Supp. Fig. S5A**).

In *C. elegans*, members of the CUT homeodomain family jointly control expression of pan-neuronal features (*27*). Two of the *C. elegans* CUT proteins, CEH-44 and CEH-48 are expressed pan-neuronally, while others are expressed ubiquitously (*27*). We engineered an epitope tag into the *Ppa-ceh-48* locus and found that epitope antibody staining provided robust pan-neuronal nuclear staining, with the only exception being no detection of *Ppa-*CEH-48 in the CAN neurons, a novelty compared to *C. elegans* (**Supp. Fig. S5B-C**).

### Conservation of the mesodermal glia

Moving beyond neurons, we examined one glial cell type whose developmental specification is particularly well understood, the GLR glia (*104*). Anatomically, the GLR glia display an interesting novelty in *P. pacificus* compared to *C. elegans*, since they generate extensive synaptic outputs (*18*). Molecularly, the *C. elegans* GLR glia uniquely co-express two terminal selectors, the Forkhead-type transcription factor LET-381 and the homeodomain protein UNC-30 (*104*); moreover, antibody staining reveals the uptake of GABA by the GLR glia is enabled by the GABA reuptake transporter SNF-11 (*72*). HCR staining shows that in *P. pacificus*, the GLRs also express the GABA reuptake transporter *snf-11* and they co-express the orthologs of the two identity regulators *let-381* and *unc-30* (**Fig. 11G**).

**Figure 11.**
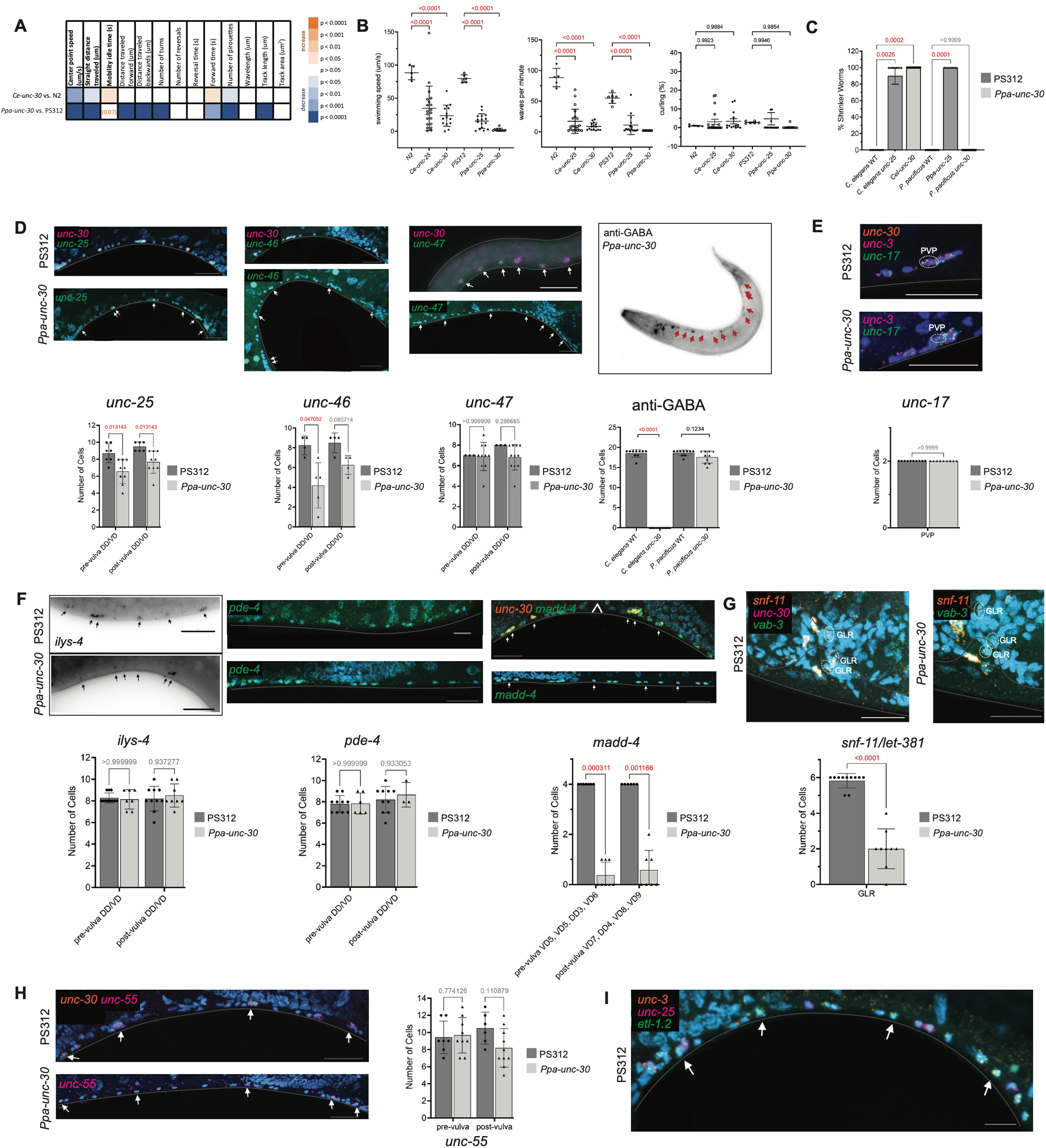
Behavioral and molecular analysis of *Ppa-unc-30* null mutation animals. **(B)** Heatmap showing comparison of locomotory characteristics between wild-type *C. elegans* (N2), *Cel-unc-30* null mutant, wild-type *P. pacificus* (PS312) and *Ppa-unc-30* null mutant. Orange indicates positive significant change, and blue indicates negative significant change. Loss of *unc-30* in both species results in a reduction of locomotion and increase in mobility idle time as expected of a ‘unc’ phenotype. **(C)** Quantitative analysis of tracked swimming of wild-type, *unc-25* null mutants, and *unc-30* null mutants of both *C. elegans* and *P. pacificus*. Loss of *unc-25* leads to reduction in swimming speed, waves per minute, and slight increase in curling in both species. Loss of *unc-30* in *C. elegans* mirrors loss of *unc-25*, while loss of *unc-30* in *P. pacificus* results in complete paralysis and inability to swim (swimming speed, waves per minute, and curling = 0). **(D)** Quantitative analysis of shrinker assays of wild-type, *unc-25* mutants, and *unc-30* mutants of both *C. elegans* and *P. pacificus*. Loss of *Cel-unc-25* and *Cel-unc-30* leads to an increase in the ‘shrinker’ phenotype in response to head touch. *Ppa-unc-30* null mutants don’t shrink. **(E)** *unc-30* weakly regulates GABAergic identity of the D-type neurons in the VNC. HCR RNA FISH against GABAergic identity markers *Ppa-unc-25* and *Ppa-unc-46* is reduced, while *Ppa-unc-47* and anti-GABA remain unchanged. Arrows indicate neurons still expressing GABAergic identity markers. **(F)** Loss of *Ppa-unc-30* does not result in the loss or reduction of *Ppa-unc-17* in the PVP neuron in the tail. **(G)** *Ppa-unc-30* does not regulate D-type terminal identity markers (HCR signal) *Ppa-ilys-4* or *Ppa-pde-4*, as indicated by lack of effect in *Ppa-unc-30* mutants. However, loss of *Ppa-unc-30* does result in the reduction of *Ppa-madd-4*. Arrowhead indicates position of the vulva. **(H)** *Ppa-unc-30* regulates expression of *Ppa-snf-11* in the mesodermal glial cells GLR. Expression of *Ppa-let-381* is unaffected. **(I)** Loss of *Ppa-unc-30* does not affect the expression of *Ppa-unc-55* in the D-type neurons of the VNC. All images shown are single z-slices or projections of >3 z-slices of adult hermaphrodites acquired with confocal microscopy. **(J)** HCR staining of *Ppa-elt-1.2* indicates that unlike in *C. elegans*, where *unc-30* and *elt-1* are co-expressed and co-operate in the D-type neurons, *Ppa-elt-1.2* is not expressed in *Ppa* D-type, but instead in *Ppa-unc-3*-positive cholinergic neurons. Scale bars indicate 20 µm. Arrows indicate retention of terminal identity marker signal and/or overlap of relevant HCR probes. All P-values calculated by t-test or one-way ANOVA.

### Knockouts of *P. pacificus* terminal selectors and subtype selectors

Moving beyond the description of expression patterns of putative regulatory factors and markers that help define cellular identities, we next set out to probe the conservation of genetic regulatory signatures on a functional level. To this end, we engineered null alleles of eight *P. pacificus* transcription factors discussed above, focusing on those transcription factors whose loss results in well-defined molecular and behavioral defects in *C. elegans*, namely *unc-86, mec-3, unc-42, unc-3, unc-30, unc-4, ceh-10* and *lim-6* (**Supp. Fig. S1** and **Methods**). As we will describe in the next sections, this genetic analysis revealed a rich set of similarities, as well as some divergences in gene function.

### Effect of loss of *Ppa-unc-86/Brn3* on neuronal differentiation

We used the CRISPR/Cas9 system to engineer a full locus deletion of the *Ppa-unc-86* POU homeobox gene (**Supp. Fig. S1**). As in *C. elegans, Ppa-unc-86* mutant worms are viable and display no obvious morphological defects. As in *C. elegans*, and consistent with its expression in touch receptor neurons, *Ppa-unc-86* mutants are touch-insensitive (**Fig. 7A**). Consistent with a recent report (*105*), we observed the same touch insensitivity in *Ppa-mec-3* mutants (**Fig. 8B**) Downstream targets of *unc-86* in the touch receptor neurons are conserved, since we found expression of *mec-3* and *eat-4/VGluT*, two markers of touch neuron differentiation (*33, 106*) to be lost in *Ppa-unc-86* mutants (**Fig. 7C**).

In the pharyngeal, serotonergic NSM neurons, all three markers tested (*tph-1, cat-1, mod-5*, anti-5HT) are strongly affected in *Ppa-unc-86* mutants (**Fig. 7D, Fig. S6**). The extent of effects is stronger than in *Cel-unc-86* mutants, where several of these markers are only completely affected upon the simultaneous removal of *unc-86’s* co-factor *ttx-3* (*41*). The serotonergic phenotype of the I1 pharyngeal neuron depends, at least in part on *Ppa-unc-86* (**Fig. 7D**). Hence, *Cel-unc-86* is apparently not sufficient to induce serotonergic identity in *C. elegans* I1, but has acquired this ability in *P. pacificus*, likely in combination with other transcription factors.

Within the anterior deirid lineage, *Cel-unc-86* promotes the fate of distinct neuron classes, including the peptidergic RMG neurons (*26*). Similarly, *Ppa-unc-86* also affects *nlp-56* neuropeptide expression in RMG (**Fig. 7E**). Apart from promoting specific fates in this lineage, *unc-86* simultaneously suppresses the execution of dopaminergic neuron fate (*29*), at least in part through the repression of dopaminergic fate regulators such as *ast-1* and *ceh-43* (*26*). Ectopic dopaminergic neurons are also evident in the postdeirid lineage of *Cel-unc-86* mutants (*29*). The same regulatory architecture appears to exist in *P. pacificus*, as evidenced by the derepression of ectopic dopaminergic fate markers in the anterior and posterior deirid region of *Ppa-unc-86* mutants (**Fig. 7F**).

We observed a *Ppa-unc-86* mutant phenotype that points to a regulatory linkage unique to *P. pacificus*. Unlike in *Cel-unc-86* mutants, *Ppa-unc-86* mutants occasionally display anti-serotonin staining in peripheral neurons along the body wall (**Supp. Fig. S6**). While we can presently not assign an identity to these neurons, we consider it most likely that normally UNC-86-dependent neurons in the body wall may transform their identity to a serotonergic neuron in the absence of *Ppa-unc-86*.

The Ppa-AIM neuron pair, which does not display the glutamatergic identity that its *C. elegans* counterpart displays, shows a divergent dependence of *unc-86*. While in *C. elegans, unc-86* controls both the glutamatergic as well as the expression of the neuropeptide encoding genes, *nlp-73* (*26, 33*), the expression of *Ppa-nlp-73*, as well as another AIM-expressed neuropeptide, *Ppa-nlp-70*, are unaffected in *Ppa-unc-86* mutant animals (**Fig. 7G**).

In the head anterior ganglion, we observe some apparent differences in *unc-86* gene function. The cholinergic identity of the anterior ganglion neurons, IL2, URA, URB, assessed by *unc-17/VAChT* expression, are controlled in *C. elegans* by *unc-86* (*41*) but their *P. pacificus* counterparts retain *unc-17/VAChT* expression in *Ppa-unc-86* mutants (**Fig. 7H**). There is, however, an effect of *Ppa-unc-86* removal on the expression of the non-subtype specific *egas* genes (**Fig. 7I**). Curiously, the expression of three genes that are expressed in a non-subtype specific manner, *Ppa-egas-4, degl-1* and *klp-6* are reduced selectively in the dorsoventral, but not lateral IL2 neuron pairs, and the other *egas* genes are more severely affected in the dorsoventral IL2s than the lateral IL2s (**Fig. 7I,J**). These observations indicate a shift of function of *Ppa-unc-86* compared to *Cel-unc-86* and support the existence of an intermediate subtype in the Ppa-IL2s.

### Effect of loss of *Ppa-unc-42* on neuronal differentiation

We used the CRISPR/Cas9 system to engineer two deletion alleles of the Prop1-type homeobox gene *Ppa-unc-42* that are predicted to disrupt the coding region of the locus (**Supp. Fig. S1**). As in *C. elegans* (*107*), these animals display uncoordinated locomotion, readily quantifiable by measuring speed and idle time (**Fig. 8A; Fig. S9**). Apart from locomotory defects, *Cel-unc-42* mutants also display defects in nociception, likely caused by differentiation defects of the ASH nociceptive neuron class (*107*). *Ppa-unc-42* is also expressed in ASH and, like *Cel-unc-42* mutants, *Ppa-unc-42* mutants display defects in the response to nose touch and octanol, a noxious chemical (**Fig. 8B**). Matching this behavioral defect, the ASH neuron class fails to express its glutamatergic phenotype (i.e. *Ppa-eat-4/VGluT* expression) in *unc-42* mutant animals (**Fig. 8C**), as observed in *C. elegans unc-42* mutants. ASH retrains amphid neuron characteristics in *Ppa-unc-42* mutants: it still fills with lipophilic dye (**Supp. Fig. S7)** and retains expression of the ciliary marker *osm-6* (**Fig. 6D**).

Locomotory defects of *Ppa-unc-42* mutants are mirrored by differentiation defects in neurons that control locomotory behavior. A group of cholinergic neurons, including the command interneurons, as well as motor neurons in the ventral ganglion, lose their cholinergic phenotype (i.e. *unc-17/VAChT* expression) in *Ppa-unc-42* mutants (**Fig. 8D**). Again, these molecular phenotypes are similar to those observed upon loss of *Cel-unc-42* (*65, 107*).

In the ventral ganglion, *Cel-unc-42* acts as a terminal selector for the neuropeptidergic AVK neuron class (*69*). As described above, peptidergic features of Ppa-AVK are conserved, and like in *C. elegans, flp-1* expression is lost in *Ppa-unc-42* mutants (**Fig. 8E**).

In *Cel-unc-42* mutants, AVK and other ventral ganglion neurons transform their identity into GABAergic neurons, apparently through the derepression of a GABA fate-inducing transcription factor (*72*). A comparable regulatory linkage appears to exist in *P. pacificus*, since we find ectopic GABA staining and *unc-25/GAD* HCR signals in the ventral ganglion of *Ppa-unc-42* mutants (**Fig. 8F**).

### Effect of loss of *Ppa-unc-3* on neuronal differentiation

We used the CRISPR/Cas9 system to engineer a complete locus deletion of the COE-ortholog *Ppa-unc-3* (**Supp. Fig. S1**). Loss of *Ppa-unc-3* results in a behavioral phenotype very similar to that of *Cel-unc-3* mutants. Mutant animals are nearly paralyzed, as expected from the loss of proper cholinergic motor neuron differentiation (**Fig. 9A; Fig. S9**). Indeed, like in *C. elegans* (*83*), the cholinergic phenotype of normally UNC-3-expressing ventral nerve cord motor neurons is lost, leaving *unc-17/VAChT* expression intact in only a few ventral nerve cord neurons (**Fig. 9B**), which we presume to be the serotonergic VC neurons. Anti-cholinergic protein staining (UNC-17 + CHA-1) also shows a strong reduction in cholinergic signal along the ventral and dorsal nerve cords in *Ppa-unc-3* mutants, while leaving – as expected – signal in the sublateral cords unaffected (**Fig. 9C**); a similar reduction in staining is observed in *Cel-unc-3* mutants (**Fig.10C**). Illustrating conservation of the breadth of *Ppa-unc-3’s* effect on motor neuron differentiation, we also find that expression of the synaptic organizer gene *madd-4* is severely reduced in *Ppa-unc-3* mutant animals (**Fig. 9D**).

In *C. elegans, unc-3* acts in a feedforward manner to regulate motor neuron-class specific regulatory factors, which in turn promote or antagonize the function of *unc-3* in individual neuron classes (*84*). This regulatory architecture of motor neuron class diversification also appears to be largely conserved, since the expression of the class-specific regulators *unc-4* (DA/VA), *bnc-1* (VA/VB), *vab-7* (DB) is affected in the respective neuron classes of *Ppa-unc-3* mutants (**Fig. 9E,F;** as expected, *unc-4* and *vab-7* remain unaffected in the normally UNC-3-negative VC neurons). There is no observed effect on *mab-9* or *unc-55*, which are co-expressed in the AS subclass (**Fig. 9F, G**).

### *Ppa-unc-4* deletion reveals conservation of cholinergic motor neuron class diversification

While the expression of *unc-4* in A-type motor neurons, as well its regulation by *unc-3* indicates conservation of gene function, we sought to experimentally confirm such conservation through the generation of *Ppa*-*unc-4* mutants. We found that these mutant animals indeed display the forward locomotory defect expected from a loss of A-type motor neuron functionality (*108*)(**Fig. 10A; Fig. S9**). The Mnx-type homeobox *ceh-12*, is expressed in VB motor neurons in *C. elegans* and repressed by *Cel-unc-4* in VA motor neurons to shape the proper synaptic wiring of the VA neurons (*86*). Its *P. pacificus* ortholog, *Ppa-ceh-12*, shows the same VB-type specific expression (**Fig. 10B**). Moreover, as in *C. elegans, Ppa-ceh-12* expression is derepressed in VA motor neurons in *Ppa-unc-4* mutants (**Fig. 10B**). However, the target gene spectrum of *unc-4* also shows some divergences. In *C. elegans*, the *unc-4* phenotype is manifested molecularly by a loss of A-type markers and gain of B-type markers instead. We find that the B-type-specific ACh receptor *acr-5*, repressed by *Cel-unc-4* in the B-type neurons (*109*), is not expressed in B-type neurons in *P. pacificus*, neither is another *C. elegans* B-type specific markers, *nlp-71*.

Apart from its classic role as a subtype selector of the A vs. B class of motor neurons, *Cel-unc-4* operates as a terminal selector in the *C. elegans* VC motor neurons (*92*). In *C. elegans* a subset of ventral cord VC neurons, VC4 and V5, are serotonergic, but acquire their serotonin not through *tph-1*-dependent synthesis, but via *mod-5/SERT-*dependent uptake of serotonin released by other neurons (*67, 110, 111*). In contrast, in *P. pacificus* all VC neurons are serotonergic, and they are capable of producing their own serotonin, producing their own serotonin via *tph-1* expression (*37*). We found that the serotonergic phenotype of all Ppa-VC neurons is diminished in *Ppa-unc-4* mutants (**Fig. 10C**). Hence, while *Cel-unc-4* can induce a serotonergic phenotype only in a subset of VC neurons in *C. elegans*, its *P. pacificus* ortholog controls the serotonergic phenotype of all VC neurons.

### *unc-30* deletion results in unexpected phenotypes

We used the CRISPR/Cas9 system to engineer a complete locus deletion of the *Ppa-unc-30* Pitx-type homeobox gene, the ortholog of the D-type motor neuron terminal selector *Cel-unc-30* (**Supp. Fig. S1**). In *C. elegans*, loss of *unc-30* results in striking differentiation defects of D-type motor neurons, including the D-type motor neuron-specific loss of expression of GABA production and usage, based on loss of expression of *unc-25/GAD* and *unc-47/VGAT*, the enzyme that synthesizes and transport GABA (*96, 97*). GABA is required for body wall muscle relaxation and hence, both *Cel-unc-30* and *Cel-unc-25* mutants display a “shrinker” phenotype, as well as swimming defects, a thrashing-like movement in liquid (*112*).

Surprisingly, *Ppa-unc-30* mutants show a distinctive locomotory behavior. They are much more severely defective in locomotion (crawling) than *Cel-unc-30* mutants and display more defective swimming behavior (**Fig. 11A-C**). Moreover, *Ppa-unc-30* mutants do not shrink in response to prodding (**Fig. 11D**). To assess whether the lack of a shrinker phenotype is reflective of a different usage of GABA in *P. pacificus*, we generated *Ppa-unc-25* mutants by CRISPR/Cas9 genome engineering and found that these animals phenocopy the shrinker phenotype of *Cel-unc-25* mutants (**Fig. 11D**). *Ppa-unc-25* also displays swimming defects comparable to those of *Cel-unc-25* mutants (**Fig. 11C**).

The molecular functions of *Ppa-unc-30* and *Cel-unc-30* as D-type motor neurons identity regulators also differ. Unlike in *C. elegans, Ppa-unc-30* does not affect GABA staining in the ventral nerve cord (**Fig. 11E**). Expression of *P. pacificus* orthologs of GABA synthesis and transport pathway genes that are regulated by *unc-30* in *C. elegans*, namely *unc-25/GAD, unc-47/VGAT* and *unc-46/LAMP*, are also only very mildly affected (*unc-25)* or not affected at all (*unc-46, unc-47)*(**Fig. 11E**) in *Ppa-unc-30* null mutants. To ensure that the differences in *unc-25/GAD* regulation are not the result of different technology used (reporter gene analysis in *C. elegans*, transcript measurement in *P. pacificus)*, we used HCR to stain for *Cel-unc-25* transcripts in wildtype *C. elegans* and *unc-30* mutants and corroborated a complete loss of transcript (**Supp.Fig. S8**).

The *P. pacificus* orthologs of two other targets of *Cel-unc-30, Ppa-ilys-4* and *Ppa-pde-4*, are expressed in Ppa-D-type motor neurons as well but also fail to be regulated by *Ppa-unc-30* (**Fig. 11F**). Expression of the *P. pacificus* ortholog of the synaptic organizer *madd-4*, another *Cel-unc-30* target is, however, strongly affected by *Ppa-unc-30* (**Fig. 11F**).

Outside of the ventral nerve cord, *Ppa-unc-30* is expressed in PVP (**Fig. 11E**), a cholinergic pair of neurons in the pre-anal ganglion as in *C. elegans* (*64, 97*). While *Ppa-unc-30* expression in PVP is conserved in *P. pacificus*, its regulatory function appears not to be. Contrasting the role of *unc-30* in controlling the cholinergic phenotype of *C. elegans* PVP (*64*), *unc-17/VAChT* expression is not affected in PVP in *Ppa-unc-30* mutants (**Fig. 11E**). Lastly, in GLR mesodermal glia, *Ppa-unc-30* function appears to be conserved: Like in *C. elegans* (*104*), *Ppa-unc-30* mutant show a decrease in *snf-11* expression in the GLRs (**Fig. 11G**).

### Essential functions of two homeobox genes point to similarities and divergences in function

The *ceh-10* Prd-type homeobox gene is required for the differentiation of the CAN neuron pair in *C. elegans*, one of only two neuron classes required for viability in *C. elegans* (*113*). Loss of CAN cell differentiation results in the formation of vacuoles due to failures to engage in osmoregulation, causing a completely penetrant L1 larval arrest (*113*). We found that a full locus deletion of *Ppa-ceh-10*, using CRISPR/Cas9 similarly results in early larval arrest with vacuoles forming throughout the body (**Fig. S10**). Expression and function of *ceh-10* therefore appears to be conserved between *C. elegans* and *P. pacificus*.

We also sought to assess the function of the LIM homeobox gene *lim-6*, which we find to display the same expression pattern in GABAergic neurons in *P. pacificus* as in *C. elegans*, as described above. The knockout of *lim-6* in *C. elegans* results in completely viable animals that display differentiation defects in the RIS, AVL and DVB GABAergic neurons in which *lim-6* is normally expressed in (*55, 72*). However, while we were able to isolate heterozygous *Ppa-lim-6(-)* animals through CRISPR/Cas9 genome engineering, no viable homozygous offspring could be identified, and dead embryos were observed on the plate. This observation indicates that *lim-6* function in *C. elegans* and *P. pacificus* has diverged to include an essential embryonic function.

## DISCUSSION

### Neuronal gene expression changes in distant nematodes

Our analysis provides a rich, panoramic view of patterns of molecular and functional evolutionary change across two nervous systems separated by over 200 million years of evolution. Our prior comparison of the molecular composition of the brain of three distinct species from the *Caenorhabditis* genus, separated by ~ 45 million years of evolution showed that (a) there are no genomic novelties in terms of terminal selector gene complement (i.e. 1:1 orthology, no gain or losses); (b) that there is pervasive conservation of neurotransmitter identity and terminal selector expression; and (c) that the main sources of novelties concerned diverged expression of neurotransmitter receptors, as well as neuropeptide-encoding genes and their receptors (*14*).

In contrast, our present comparative analysis of *C. elegans* with *P. pacificus* revealed divergences on all three levels. While terminal selectors, particularly of the homeodomain-type are deeply conserved, reaching across all animal phyla, there are a few duplications event that either occurred in *C. elegans* or *P. pacificus* or independently in both. A comparison of the key functional feature of neuronal cell types, neurotransmitter identity, reveals broad patterns of conservation between *C. elegans* and *P. pacificus*, with several novelties occurring throughout the nervous system. For example, the peripheral BDU interneurons, peptidergic in *C. elegans*, are cholinergic in *P. pacificus*. The RIP head interneuron is cholinergic in *C. elegans*, but serotonergic in *P. pacificus*. This change in neurotransmitter identity is concordant with striking synaptic connectivity differences between *C. elegans* and *P. pacificus* RIP (*18*), which may underlie the altered feeding behavior of both nematodes. Three neuron classes show an altered glutamatergic identity: The AIM inter- and OLQ sensory neuron classes are glutamatergic only in *C. elegans*, whereas the RMGR motor neuron is glutamatergic only in *P. pacificus*. Intriguingly, this glutamatergic identity seems to be lateralized, i.e. employed only by the right, but not the left RMG neuron, adding to the very rare examples of neuronal laterality in nematodes. We furthermore note that the change in glutamatergic identity of the AIM neuron class is also concordant with alterations in synaptic connectivity; glutamatergic Cel-AIM stands out in the number Cel-specific synapses that the neuron generates, compared to its *P. pacificus* homolog (*18*). On the other hand, like the previously described dopaminergic, octopaminergic and tyraminergic neuron identities (*37*), we find sites of GABAergic neurotransmitter synthesis to be completely conserved among the two nematodes.

On the level of transcription factors, we observed broad patterns of conservation throughout the nervous system, but again with several notable differences. There are several cases in which a *C. elegans* terminal selector is expressed in additional neuron classes in *P. pacificus* (e.g. *che-1* in AFD and *unc-42* in amphid sensory neuron). Conversely, we observed several cases, in which *C. elegans* terminal selectors are not expressed in the same neuron class in *P. pacificus* (e.g. *elt-1* in D-type motor neurons, *mls-2* in AIM, *hlh-3* in VC & HSN neurons). In most cases, these differences are paralleled by notable alterations in phenotypic features of these neurons. For example, the AIM neurons alter their glutamatergic neurotransmitter identity as well as their serotonin-uptake function and the IL2 neurons show differences in their subtype diversification program (summarized in **Table 1**).

There are notable changes in the expression of subtype selectors that speak to the evolutionary lability of neuronal subtype diversification programs. The subtype selector *unc-39* diversifies the IL2 class in *C. elegans* by promoting the expression of a IL2L/R features and inhibiting IL2D/V features (*45*), but its *P. pacificus* orthologue is not expressed in the *P. pacificus* IL2 neurons. Consequently, the subtype diversification of IL2 neurons, based on terminal marker expression, appears altered in *P. pacificus*, with several subtype-specific *C. elegans* genes either not being expressed in Ppa-IL2s at all, or expressed in all IL2 neurons. Conversely, the left and right ASE neurons, which are diversified, in part, by the ASEL-specific *lim-6* gene in *C. elegans* (*56*), show a bilaterally symmetric expression of *lim-6* in *P. pacificus*. The ASEL/R neurons are lateralized in *P. pacificus* as well (*54*) but must use different regulatory factors to achieve such lateralization. The outer labial neurons, which are so distinctive in *C. elegans* that they have been given different names (OLQ for the dorsal/ventral pairs of OL neurons; OLL for the lateral pair)(*47*), have altered the manner by which they are distinctive in *P. pacificus*. Expression of a TRP-channel (*ocr-4*) expanded to all outer labial neurons while their glutamatergic identity became restricted to only the quadrant Ppa*-*outer labial neurons.

There are also changes in the expression of transcription factors that do not fall into the terminal selector category, with possibly profound consequences on neuronal function. The *unc-30* homeobox gene does not affect the overall differentiation of the ASG neurons in *C. elegans* but is required in ASG to control some presently ill-defined immunity-related function of Cel-ASG (*52*). *P. pacificus* does not express *Ppa-unc-30* in ASG, perhaps hinting towards a divergence of ASG’s involvement in immunity control in *P. pacificus*. The *unc-3* transcription factor, a bona fide terminal selector of cholinergic motor neurons in both *C. elegans* and *P. pacificus*, has a complex function in the *C. elegans* ASI neurons, activating a subset of ASI-expressed genes, including the TGFβ protein *daf-7*, not affecting other ASI-expressed genes, but also actively repressing the expression of non-ASI expressed genes in ASI (*49, 50*). In *P. pacificus*, the terminal selector function of *unc-3* is conserved in the cholinergic ventral nerve cord motor neurons, but *Ppa-unc-3* is not expressed in ASI. The absence of ASI expression is paralleled by a notable divergence in the deployment of TGFβ in dauer formation between the two species. In *C. elegans, unc-3*-mediated control of TGFβ-like *daf-7* gene expression in ASI is required for dauer formation (*50*), while in *P. pacificus, daf-7*-encoding genes have multiplied, but apparently none are involved in dauer formation (*51*). Together, these findings indicate that the function of the ASI sensory neurons may have significantly diverged among these two nematode species, a likely reflection of the different lifestyle and ecological niches that they occupy.

### Similarities and divergences in terminal selector function

#### Similarities

The feature that sets our study most apart from the presently widely used comparative molecular profiling approaches is that ours involves comparative functional analyses based on genetic loss of function data that includes single neuron and single target gene resolution, as well as behavioral readouts. We extract two broad conclusions from our functional analysis. On the one hand, we observed many cases of similarities of *Cel-*and *Ppa-* terminal selector behavioral mutant phenotypes. *Cel-*and *Ppa-unc-42* mutant animals display similar locomotory and chemosensory defects. *Cel-* and *Ppa-unc-3* animals are both severely uncoordinated (Unc), *Cel-*and *Ppa-unc-4* animals share a characteristic forward Unc phenotype. *Cel-*and *Ppa-unc-86* animals are mechanosensory defective, and *Cel-*and *Ppa-ceh-10* both control excretory system function.

In many neuron classes, we also observe the same effects of orthologous *Cel-* and *Ppa-*terminal selectors on downstream target genes (summarized in **Table S3**). For example, *Ppa-unc-86* affects *mec* gene expression in touch neurons, serotonergic identity in the NSM neurons and glutamatergic, cholinergic or peptidergic identity in other neurons. *unc-42* and *unc-3* affect cholinergic identity in both species, i.e. *unc-17/VAChT* expression in similar neuron classes and *unc-42* affects neuropeptide expression in a peptidergic hub neuron (AVK) in both species.

Moreover, regulatory linkages between transcription factors are conserved. For example, *unc-3* regulates the expression of several motor neuron class-selector genes that diversify *unc-3*-dependent motor neuron differentiation in the ventral nerve cord (all, except for *mab-9*). The regulatory linkage of *unc-86* and *unc-42* to alternative differentiation programs is also conserved. Both *Cel-* and *Ppa-unc-42* do not only promote peptidergic and cholinergic fate in head ventral ganglia but also repress the execution of alternative GABAergic differentiation programs. Similarly, both *Cel-* and *Ppa-unc-86* mutants display conversion of several neuron classes into dopaminergic identities, indicating that *unc-86* does not only turn on differentiation gene batteries, but actively suppresses alternative differentiation programs, likely via antagonizing the expression and/or function of terminal selectors for such alternative differentiation programs. Apart from these conserved regulatory architectures, we can also predict a novel regulatory linkage: In *Ppa-unc-86* mutants, midbody neurons aberrantly express serotonergic identity, indicating that *Ppa-unc-86* may not only promote the differentiation of these neurons, but also repress alternative differentiation program(s).

#### Divergences

We discovered a range of divergences in selector gene function, on several different levels. The distinct viability phenotypes of the LIM homeobox gene *lim-6* represent the most dramatic phenotypic difference and argues for the recruitment of *Ppa-lim-6* into embryonic patterning events that are absent in *C. elegans*. More commonly, our mutant analysis discovered that within a given cell type, putative terminal selector losses result in distinctive effects on downstream target expression. For example, while *Cel-* and *Ppa-unc-86* expression is conserved in a subset of sensory neurons in the anterior ganglion, loss of *unc-86* does not seem to affect their cholinergic differentiation as obviously in *P. pacificus* as it does in *C. elegans*. Vice versa, the loss of *unc-86* has a stronger effect on the serotonergic NSM neurons in *P. pacificus* as compared to *C. elegans*. The reason for these differences may relate to observations that many years of null mutant analysis of putative terminal selector in *C. elegans* have revealed: the individual loss of two co-expressed transcription factors alone produces modest or no phenotype on the differentiation of the neuron, but the joint removal produces dramatic synergies (*24, 26, 41*). One example is the NSM neuron class, in which loss of *unc-86* affects strongly *tph-1* expression but has only mild or no effects on *mod-5/SERT* and *cat-1/VMAT* expression and serotonin staining in *C. elegans* (*41*); yet all these genes are completely turned off upon joint removal of *C. elegans ttx-3*, whose removal alone has no effect on the expression of either of these genes (*41*). In *P. pacificus*, which retains co-expression of *unc-86* and *ttx-3* in NSM, the expression effects on the same features are much stronger in *unc-86* null mutants. Therefore, it is conceivable that the impact of *unc-86* null mutants in neuron classes can be masked by cooperating co-factors and that the extent of such masking may be highly evolvable, i.e. more apparent in some species compared to others. Such masking or “buffering” phenomenon may simply be due to the presence of multiple, independent binding sites for each terminal selector of a neuron class, with the importance of each site being variable across phylogeny. In other words, changes in the *cis*-regulatory architecture of target genes may determine the species-specific importance of individual members of a regulatory signature.

Gains or losses of expressions of members of a transcription factor combination may also explain why terminal selectors appear to gain the ability to turn on specific target genes. For example, *unc-4* is apparently not alone able to turn on serotonin production in the complete set of VC neurons in *C. elegans* but has acquired this ability in *P. pacificus*, either through the gain of a cooperating factor or through the gain of binding site(s) in genes required for serotonin production. A conceptually similar observation has recently been made in *C. angaria*, where *unc-4* has gained the ability to control *mod-5* expression via gain of a *cis*-regulatory element (*114*).

The striking case of the Pitx-type *unc-30* gene may reveal changes other than the *cis-*regulatory architecture of target genes. While the expression of *unc-30* in D-type motor neurons is conserved in both species, *Ppa-unc-30* is not required to turn on, at least to the same extent, orthologs of a host of target genes of *Cel-unc-30*. Yet the overall behavioral phenotype of *unc-30* mutants is much stronger in *P. pacificus* than it is in *C. elegans*. This can be interpreted in several distinct ways: (a) D-type motor neurons may have acquired an *unc-30-*dependent (and *unc-25/GAD-* independent) function in locomotory control in *P. pacificus* that is not apparent in *C. elegans*. (b) D-type motor neurons may have the same properties in *P. pacificus* and *C. elegans*, but in *C. elegans* the full extent of these locomotory functions may not be under control of *unc-30*. (c) *Ppa-unc-30* acts in neurons other than the D-type motor neurons to control locomotory functions. Like in *C. elegans, Ppa-unc-30* is expressed in the AVJ and PVP neurons; while these neurons have no known prominent role in locomotory behavior, they may have acquired such functions in *P. pacificus*. In either case, the striking distinctiveness of behavioral defects upon *unc-30* removal in these two nematodes species illustrate the range of scenarios by which gene and cell function may diverge over evolutionary timespans.

#### Developmental systems drift (DSD)

DSD is a concept in which homologous characters with seemingly similar morphology and function are driven by evolutionary divergent pathways (*115, 116*). Shifts in the importance of individual terminal selectors within a terminal selector combination that defines homologous neurons, discussed above, appear to be good examples of DSD. Taking this idea a step further, it is possible that homologous and functionally equivalent cell types in the two nematode species may even be specified by distinct combinations of terminal selectors. However, we note that in all cases in which we observed changes in *C. elegans* terminal selector expression in *P. pacificus*, we also observed changes in effector genes, i.e. genes that determine functional properties of a neuron, such as its neurotransmitter identity (summarized in **Table 1**). While neurons that change their neurotransmitter phenotype in two different species can still be classified as homologous based on a number of other identity traits (position, morphology, synaptic connectivity, lineage), such changes should perhaps not be considered as mere drift but are more likely to have functionally relevant consequences on the behavior of an animal. Further work in increasingly evolutionarily divergent nematode species - with correspondingly distinct patterns of behavior - would better establish the extent to which the changes we observe are the result of evolutionary drift or have been selected for.

#### Conclusions

Taken together, we extract several themes from our analysis. First, patterns of conservation and novelty in gene expression and regulation are found throughout the nervous system. This is consistent with our analysis within the *Caenorhabditis* genus, as well as with our analysis of synaptic connectivity changes in the *C. elegans* versus *P. pacificus* brain. Nevertheless, it appears that changes within the sensory periphery appear to be more widespread. Second, novelties are generated by the gain and loss of expression of terminal selectors. Third, neuronal subtype diversification appears to be particularly labile and subject to evolutionary change. Fourth, the extent and mechanism by which transcription factors control neuronal differentiation appears highly evolvable, but such changes may not necessarily always have functional consequences due to the buffered nature of neuronal differentiation programs, in which loss of individual factors in a regulatory signature may have limited effects on neuronal differentiation programs.

## MATERIAL AND METHODS

### Mutant strains and transgenes and CRISPR/Cas9 genome editing

All deletion mutant alleles, as shown in **Supp. Fig. S1**, (except *Ppa-unc-42* alleles) used in this study were generated using CRISPR/Cas9 genome engineering using two guides and a repair template designed to induce homology-directed repair and deletion of the entire gene coding region. *Ppa-unc-42(ot5021)* is a smaller deletion early in the locus that leads to a frameshift and early stop. FLAG-tagged alleles were also generated using CRISPR/Cas9 genome engineering, with one guide and a repair template designed to insert two copies of a FLAG epitope, with the exception of *unc-86*, which received a single FLAG epitope. For two loci, *unc-3* and *unc-42*, strains were also generated with two copies of an ALFA epitope, using the same guide. For every epitope-tagged gene at least two independent strains were generated and examined; for a given gene, no differences were observed in the pattern of expression in different strains. Guides, repair template, and final loci sequences are provided in **Table S4**.

### Immunocytochemistry

Anti-Cel-UNC-86 was performed as previously described (*30*) using a modified ‘Finney-Ruvkun’-style fixation and permeabilization (*117*). No staining was observed in *Ppa-unc-86* deletion mutants. Staining matched that seen with *Ppa-unc-86-*FLAG epitope-tagged strains (Supplemental figure); this method, however, had an inferior signal-to-background staining. Anti-FLAG epitope staining was also performed using the modified ‘Finney-Ruvkun’ method, but sometimes with the addition of heat induced antigen retrieval (HIAR)(*118*). HIAR was essential for UNC-86::1xFLAG staining, and greatly improved UNC-3::2xFLAG staining but was neither required nor desireable for UNC-42::2xFLAG or UNC-30::2xFLAG staining. Two different primary anti-FLAG antibodies were used: most staining was performed with a Mouse anti-FLAG monoclonal (Sigma F1804 or Invitrogen MA1-91878); UNC-30::2xFLAG staining in particular was greatly improved by using a Rat anti-FLAG (Novus Biologicals NBP106712SS). Typically, a red fluorophore-conjugated secondary antibody was used. Anti-ALFA epitope staining was performed in doubly-labeled strains by the same method, using a fluorophore-conjugated anti-ALFA nanobody (catalog no). Anti-serotonin staining was performed as previously described (*37*). Anti-ALFA epitope staining with the nanobody was also performed with anti-serotonin staining. Anti-GABA immunostaining was performed as previously described (14). Worms were fixed in glutaraldehyde and stained successively with primary and a fluorophore-conjugated secondary antibody. Worms were then kept in buffer at 4 C until imaged.

### smFISH and RNA-FISH Hybridization Chain Reaction (HCR)

smFISH was performed as previously described in Wormbook, with no optimization made for *P. pacificus* except for longer digestion time. HCR was performed as previously described by us for *P. pacificus* (*94*). Worms were fixed in paraformaldehyde, then incubated with probes for hybridization to RNA transcripts. For smFISH, these probes were conjugated with fluorophores, and for HCR, a second incubation was required to hybridize fluorescent conjugated hairpins to the probes and amplify signal. Worms were kept at 4C in ProLong Antifade Buffer until imaged (signal was retained for up to 4 months after hybridization, but for best results microscopy was performed within the same week). See **Table S5** for more detail on probes used. Probe sequences available upon request.

### Dye-filling

Dye-filling was performed in low-salt concentration for 4 hours at 1:125 concentration and rocked continuously with vigorous shaking every 30 minutes (*119*).

### Mutant analysis scoring and statistics

For HCR experiments, expression was analyzed by looking for presence of RNA-FISH puncta in each neuron (identified by DAPI signal, morphological position, and overlapping reporters of neuronal identity). At least 2 puncta near DAPI signal were required to confirm expression, and each cell was counted as “1”, so that bilaterally symmetric pairs were “2”, and so on. To determine statistical significance between groups, a t-test was performed, and the resulting P-values are shown. For immunostaining experiments, scoring and analysis was similarly carried out, with the exception of the signal being nuclear for anti-FLAG and anti-ALFA staining.

### Microscopy and Image Processing

For imaging immunofluorescence preparations, 5µl of worms in PBSTx buffer were mounted on agarose pads and covered by a square coverslip. Images were acquired on a compound upright fluorescence microscope or laser scanning confocal microsope (either a Zeiss LSM980 or Nikon A5). Single images and image stacks were examined and processed with Zeiss ZEN software, Nikon NIS-Elements, and/or FIJI. For imaging of RNA-FISH experiments, 2 µl of worms were mounted on a square coverslip and covered with a smaller round coverslip, which was then placed on a glass slide with a silicone isolator. Images were acquired at 40x using confocal laser microscopy (LSM980) or compound light microscopy and processed using Zeiss Zen software. For representative images containing more than one signal, colors shown correlate with relevant fluorophores (with the exception of AF514 and AF594, which are both represented as yellow), and for images stained with only one fluorophore, images were converted to inverted grayscale. Images shown are maximum intensity projections of a few (1-5) slices or single slices; some immunofluorescence images include montages.

### Tracking, locomotion analysis and behavioral assays

For analysis of locomotion, 5-10 worms were placed on an agar plate (without food) and 5-minute videos were acquired using the MultiWorm Tracker by MBF Biosciences. Automated tracking and editing were carried out by MBF software WormLab (*120*). For swimming analysis, the same workflow was applied for 10-minute videos. Quantitative locomotory characteristics such as speed, distance traveled, and pausing time of each group were analyzed using unpaired t-tests to determine p value and statistical significance. Full graphs of locomotory characteristics can be found in **Supp. Fig. S9**.

Behavioral assays including nociceptive assays, touch assays, shrinker assays, and paralysis assays were performed as previously described (*121*). At least 10 worms were assayed 3 times each with the relevant behavioral stimulus (i.e. touch) and an average of all 3 trials was recorded. These data were then represented as response indexes, with a score of 1.0 indicating a 100% response rate, which has been previously described (*122*).

## ACKNOWLEGEMENTS

We thank…. and XYZ for comments on the manuscript. We thank USD undergrad students Hayley Lee, Madeleine Neff, Kaya Patel, Sidney Tookes, Sydney Wong for contributions to characterizing epitope expression patterns for several genes.

