## Supplementary material for "Terminal selector and subtype selector function across 200 million years of nematode evolution": Supp

### SUPPLEMENTARY FIGURES

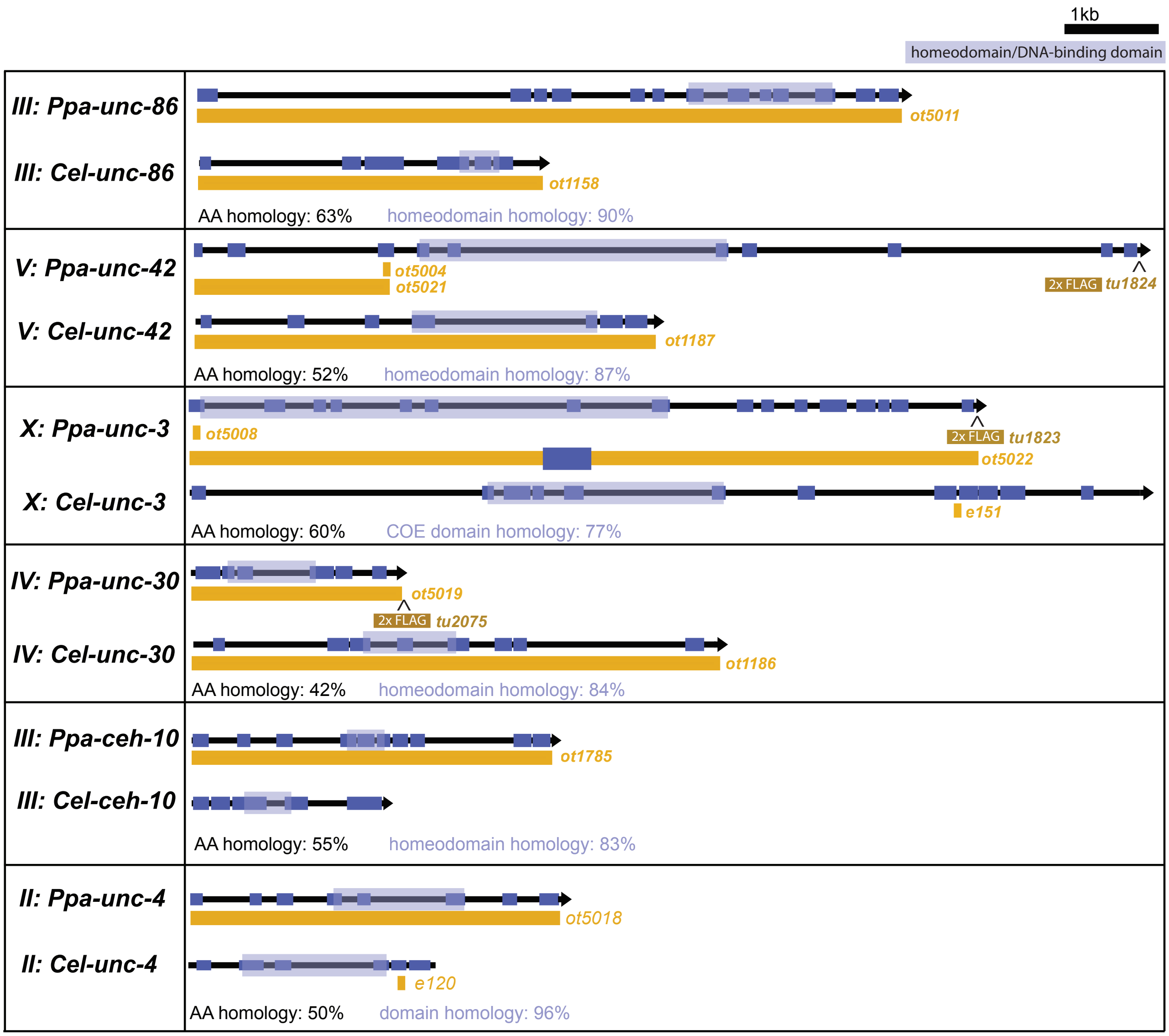


**Figure S1. Epitope-tagged and mutant alleles.** Genomic loci for FLAG-tagged, ALPA-tagged, and/or deletion mutant alleles for the investigated terminal and subtype selectors.

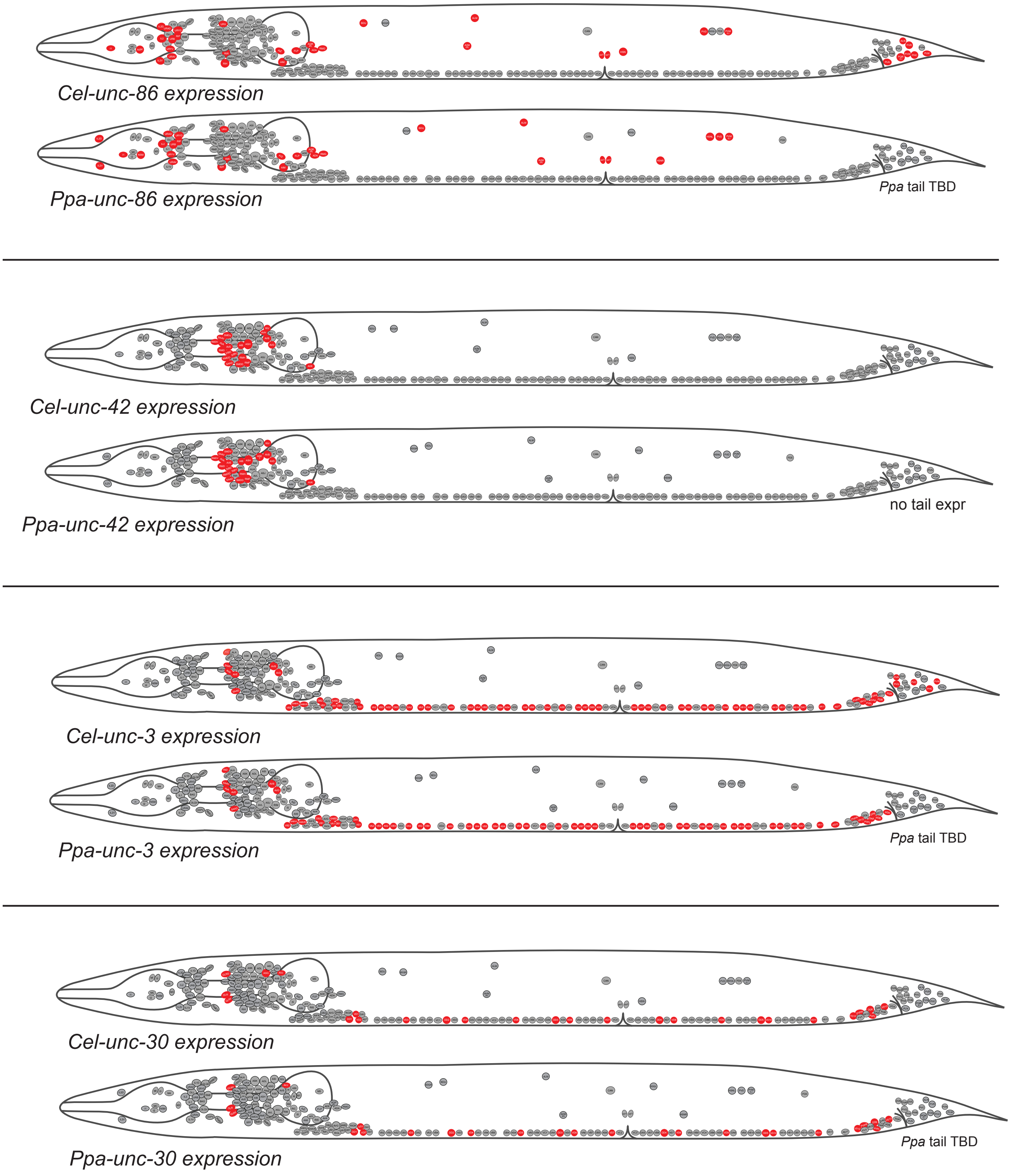


**Figure S2. Comparisons of terminal selector expression patterns in *Cel* and *Ppa*.** Summaries of expression patterns of four *C. elegans* terminal selectors compared with their *P. pacificus* orthologs as determined by antibody staining (anti-Cel-UNC-86 or anti-FLAG), shown on schematics of nuclear positions. Some prominent differences in position in *Ppa* are indicated, particularly in the body wall. (Although there are some differences in nuclear positions in the head, mostly the *Cel* positioning is maintained in the *Ppa* head schematics.) Because of complex apparent differences in the *Cel* vs *Ppa* post-anal tail, patterns of expression there will be presented in a future report (indicated as '*Ppa* tail TBD' in the schematics). A few non-neuronal nuclei are shown: uv1 cells in the vulval region, and in the *unc-30* schematics, GLR glial cells in the head.

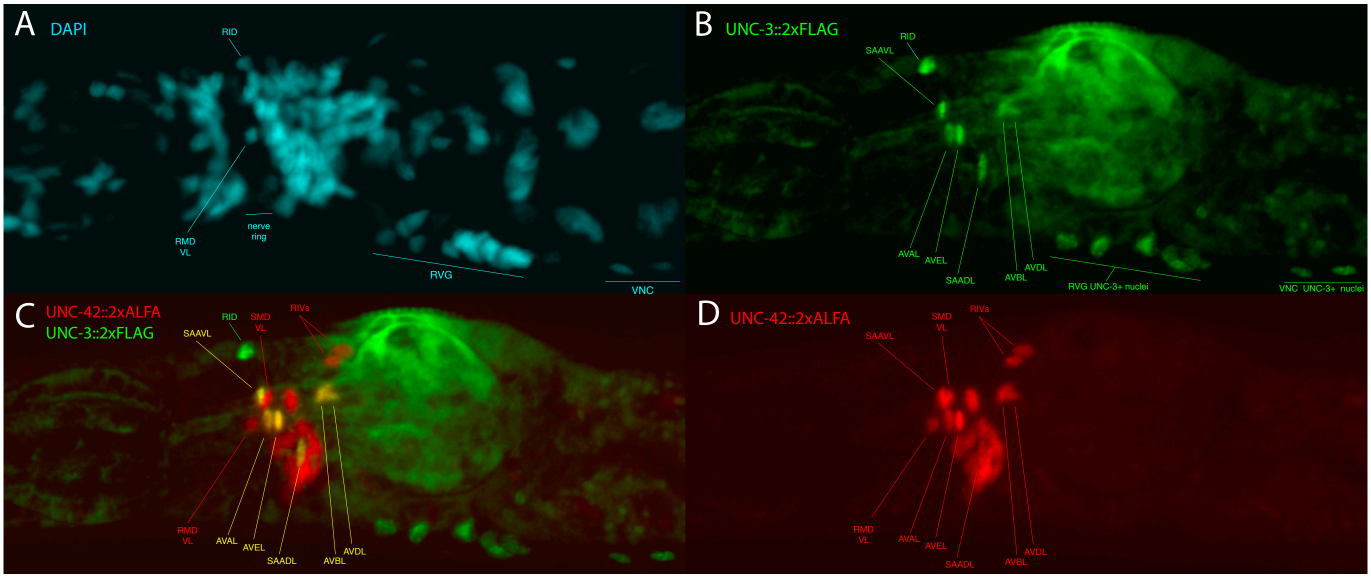


**Figure S3. Double labeling with two different epitope tags reveals co-expresson of UNC-3 and UNC-42 in command interneurons.** Anterior is to the left in all images (same focal plane, adult hermaphrodite). UNC-3::ALFA & UNC-42::2xFLAG doubly-labeled strain. (A) DAPI stained nuclei show locations of major landmarks such as the nerve ring, devoid of nuclei, and clusters such as the RVG and beginning of the VNC. Two nuclei in distinctive locations near to or overlying the nerve ring, the left side RMDV and unpaired dorsal RID, are indicated. (B) UNC-3::2xFLAG revealed by anti-FLAG staining; identified nuclei as indicated. (C) Double-labeling with UNC-3::2xFLAG and UNC-42::2xALFA; nuclei expressing both proteins are yellow, as indicated, including those of cells known in *C. elegans* as 'command interneurons.' (D) UNC-42::2xALFA revealed by red-labeled anti-ALFA nanobody staining.

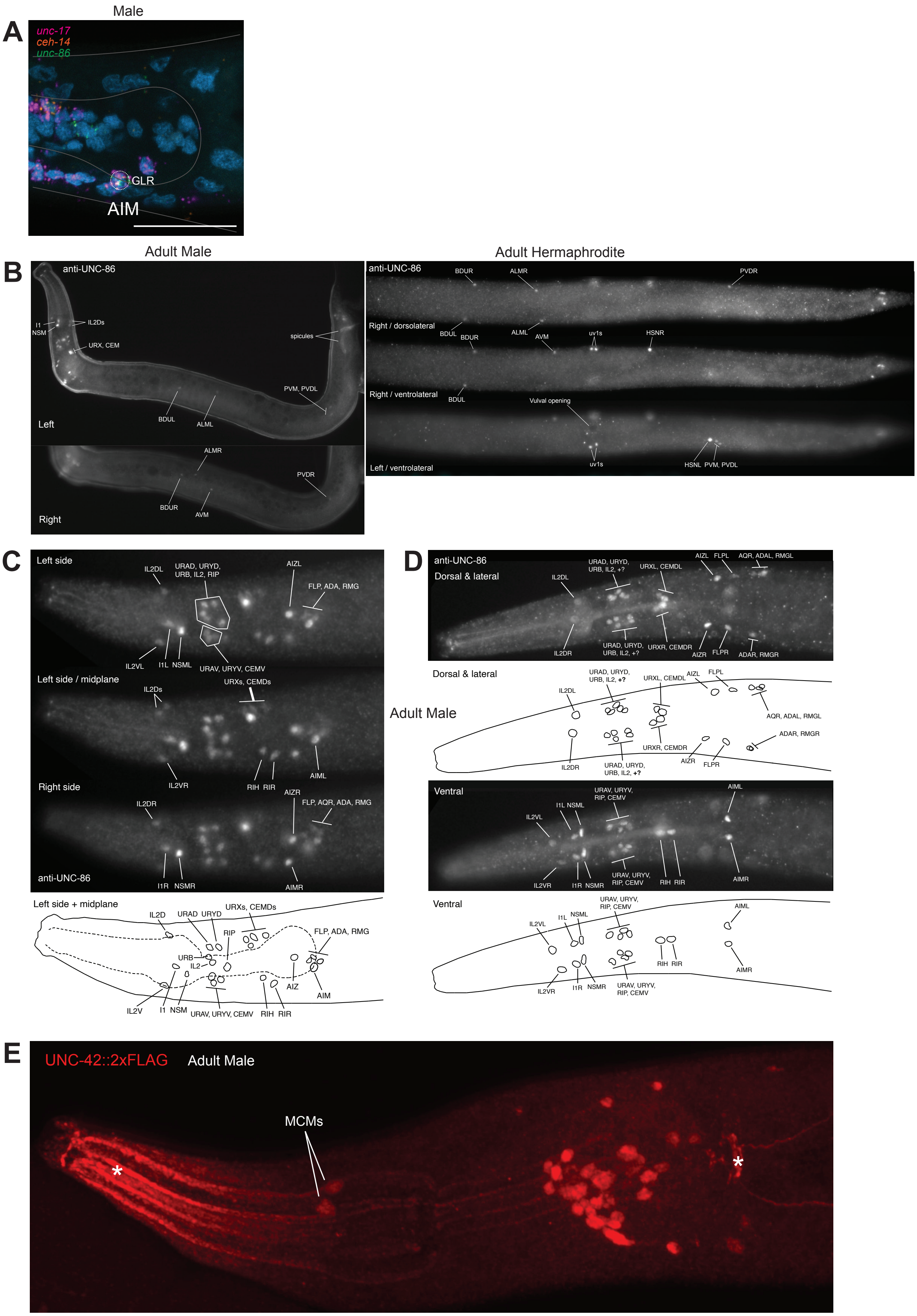

Male
A
unc-17
ceh-14
unc-86
GLR
AIM
Adult Male
Adult Hermaphrodite
B
anti-UNC-86
I1
NSM
IL2Ds
URX, CEM
spicules
PVM, PVDL
Left
BDUL
ALML
ALMR
PVDR
Right
BDUR
AVM
Right / dorsolateral
Right / ventrolateral
Left / ventrolateral
uv1s
HSNR
Vulval opening
HSNL PVM, PVDL
C
Left side
Left side / midplane
Right side
anti-UNC-86
Left side + midplane
D
Dorsal & lateral
Ventral
Adult Male
E
UNC-42::2xFLAG
Adult Male
MCMs

**Figure S4. Male-specific marker and terminal selector expression in shared and male-specific neurons.** Anterior is to the left in all images. For all lateral views, ventral is down. (**A**) Male-specific expression (via HCR) of cholinergic marker *unc-17* (magenta) in AIM, as is also observed in male *C. elegans.* AIM co-expresses m*unc-86* (green) and *ceh-14* (orange). **(B – D)** Anti-*Cel*-UNC-86 staining in *P. pacificus.* **(B)** Left panels – left and right sides of adult male, showing absence of HSNs. On the left side, a few representative head neurons are indicated. Neuronal nuclei shared with hermaphrodite can be seen, as indicated. In the tail, the auto-fluorescent male spicules are indicated. Right panels –dorsoventral/slightly lateral views of adult hermaphrodite body for contrast, with 3 focal planes showing (top to bottom) dorsal-right nuclei, ventral right, and ventral left planes with UNC-86-positive nuclei as indicated. Hermaphrodite-specific nuclei here include uv1 cells near the vulva and HSNs, located asymmetrically on the anterior-posterior axis. **(C-D)** UNC-86-positive nuclei in adult male head. **(C)** Lateral views of 3 focal plates, left side (top panel) to right side (bottom panel), with nuclei identified as indicated. All nuclei are the same as seen in hermaphrodites except for the male-specific CEM neurons: CEMVs located in the ventral anterior ganglion (anterior to the nerve ring), and CEMDs dorsally, posterior to the nerve ring. A schematic in the bottom panel identifies the nuclei in the left and midplanes, with the pharynx location outlined with dashed lines. **(D)** Dorso-ventral views of two montaged focal planes, each with a schematic below, showing an UNC-86 staining variation seen in several adult male heads. Top panels – dorsal and lateral (mid-depth) planes. In the dorsal anterior ganglion, an additional UNC-86-positive nucleus is observed on both left and right sides (noted by '+?'). A bilateral pair of CEMDs is seen posterior to the nerve right. Bottom panels – ventral plane. As in C, the same UNC-86-positive nuclei as in the hermaphrodite are seen with the addition of a bilateral pair of CEMVs in the ventral anterior ganglion. (E) Roughly lateral view of adult male head (strain RS3944), MaxIP of several focal planes to show the bilateral pair of UNC-42::2xFLAG-positive MCM neurons, as indicated. Asterisks indicate non-specific junctional staining in the pharynx (anterior) and pharyngeo-intestinal junction (posterior) seen in wildtype *P. pacificus* worms with this anti-FLAG primary antibody.

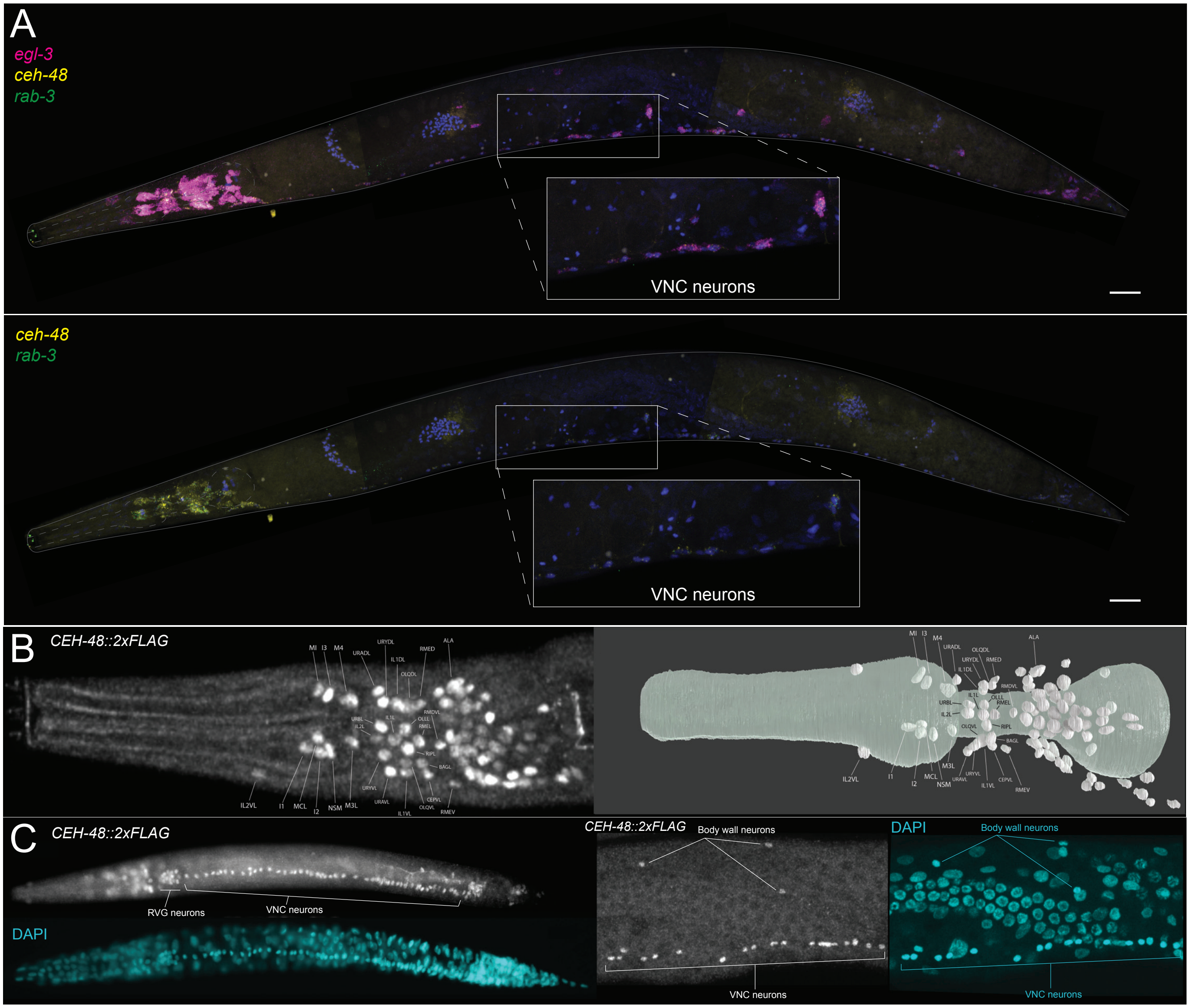


**Figure S5. Expression patterns of panneuronal identity markers *ceh-48*, *rab-3*, and *egl-3* in *P. pacificus*.**
**(A – B)** Anti-FLAG antibody staining on *P. pacificus* strain, *ceh-48(tu1612[unc-30::2xFLAG])* in which the endogenous *unc-3* locus has been tagged with a 2xFLAG epitope tag. (A) (Left) Adult hermaphrodite head lateral view with neuronal nuclei identified as indicated. (Right) Neuronal map generated from EM reconstruction of *P. pacificus* head neurons as seen in Cook et al. 2022. (B) (Left) Whole worm, ventral side, showing retrovesicular ganglion (RVG) and Ventral Nerve Cord (VNC) neurons. Anti-FLAG staining (top) lines up well with DAPI staining (bottom), indicating that CEH-48::2xFLAG staining captures all neurons in these regions. (Right) Close-up view of body wall neurons and VNC neurons also showing likeness to DAPI staining of neuronal nuclei in these regions. **(C)** Adult hermaphrodite stained with HCR RNA-FISH against *Ppa-ceh-48*, *Ppa-egl-3*, and *Ppa-rab-3* in colors as indicated. Inset shows close-up view of VNC neurons. HCR probe against *Ppa-egl-3* is bright and comprehensively captures neuronal nuclei. **(D)** Same worm showing only *Ppa-ceh-48* and *Ppa-rab-3* HCR probes, which do not clearly capture all neuronal nuclei. Scale bar 20 µm.

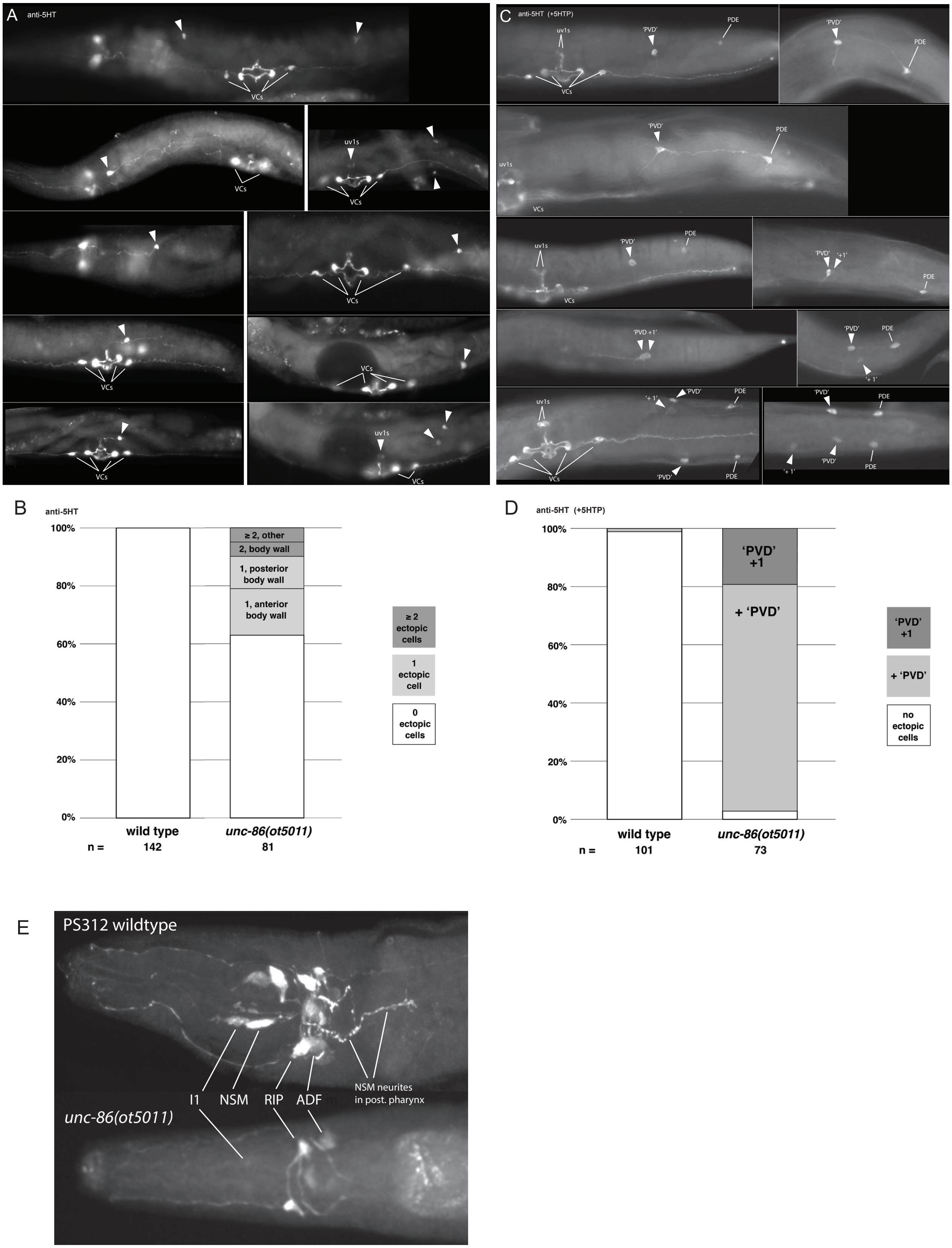


**Figure S6. Phenotypes of unc-86 mutants: mis-specific body wall neurons and loss of serotonergic identity in NSM neurons.** **(A, C, E)** Anti-5HT staining. Ectopic stained somas (A, C) are indicated with arrowheads; somas that are normally stained are indicated with lines. **(A)** Examples of mis-specified body wall neurons in *unc-*

*86(ot5009)* seen as 5HT-positive cells, including some with aberrant neurites. In wild type, serotonin-positive cells are never observed in the body wall. VC neurons in the VNC appear normal. **(B)** Quantification of ectopic 5HT-positive somas in wildtype and *unc-86(ot5011)* adult hermaphrodites. Overall, almost 40% of worms have at least one ectopic 5HT-positive soma, mostly in the body wall, about equally found in the anterior body (from pharyngeo-intestinal junction to vulva) or posterior body (vulva to anus). Occasionally, uv1s were 5HT-positive (as seen in A, bottom right panel). The 'n' indicates number of adult worms scored. **(C)** Examples of mis-specified body wall neurons in wildtype and *unc-86(ot5011)* seen as 5HT-positive cells after treatment with 5HTP, which normally makes dopaminergic neurons 5HT-positive; uv1 cells and PDE neurons in *P. pacificus* are normally stained after such treatment (Loer et al., 2026). **(D)** Quantification of ectopic 5HT-positive somas near the typical location of the PVD neuron in wildtype and *unc-86(ot5011)* adult hermaphrodites treated with 5HTP. In this preparation, a single very faint presumptive PVD soma was observed in one wild type worm. The 'n' indicates number of adult worm sides scored; sides were scored only if both PDEs and uv1s (expected to stain) could be seen on a side. **(E)** Anti-5HT staining in representative wildtype and *unc-86(ot5011)* adult hermaphrodite heads, dorsal-ventral view. Neurons as indicated on one side of each head. NSMs were never observed stained *unc-86(ot5011)* heads (0/40 heads); both somas and neurites were missing. Staining of all other head cells was normal, although possibly reduced in I1 neurons, the staining of which is weaker and more variable in wildtype.

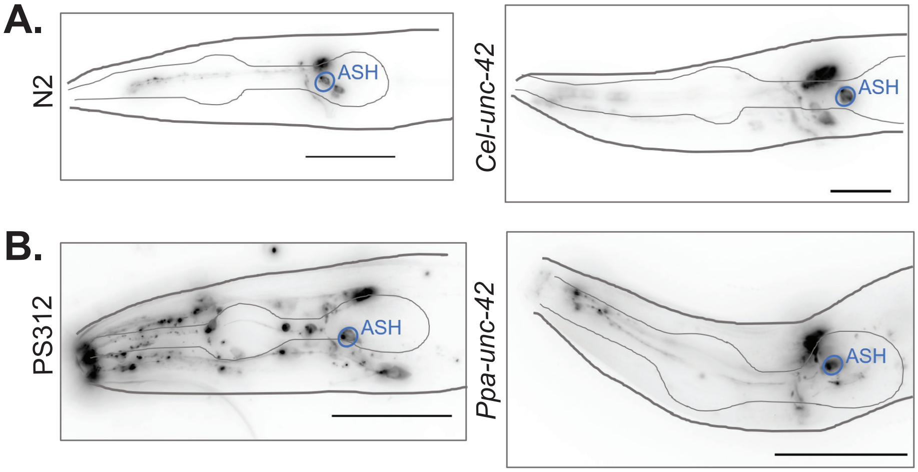


**Figure S7 Amphid sensory neurons in both *C. elegans* and *P. pacificus* fill normally with dye in *unc-42* mutants.** Anterior is to the left in all images. Heads of wildtype and *unc-42* mutant worms treated with externally applied lipophilic dye. **(A)** *C. elegans* ASH neurons fill with dye in *Cel-unc-42* mutants, like they do in wildtype (N2). **(B)** Like in *C. elegans, P. pacificus* ASH neurons fill with dye in *Ppa-unc-42* mutants, like they do in wildtype (PS312). Scale bars: 20 µm.

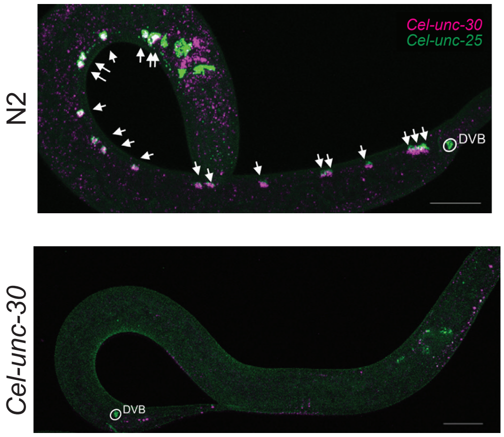


**Figure S8. Ventral nerve cord D-type motor neurons lose GABAergic identity in *Cel-unc-30* mutants.** RNA-FISH HCR in *C. elegans* with *unc-25* (green) and *unc-30* (magenta) probes. (Top panel) Wildtype (N2) shows co-expression of *unc-25* and *unc-30* genes in D-type GABAergic motor neurons (arrows). A non-VNC GABAergic neuron, DVB, expressing *unc-25* in the tail dorsorectal ganglion is circled. (Bottom) *Cel-unc-30* mutant worm showing absence of *unc-25* expression in the VNC. Expression of *unc-25* in the tail DVB neuron (circled) is normal. Scale bars: 20 µm.

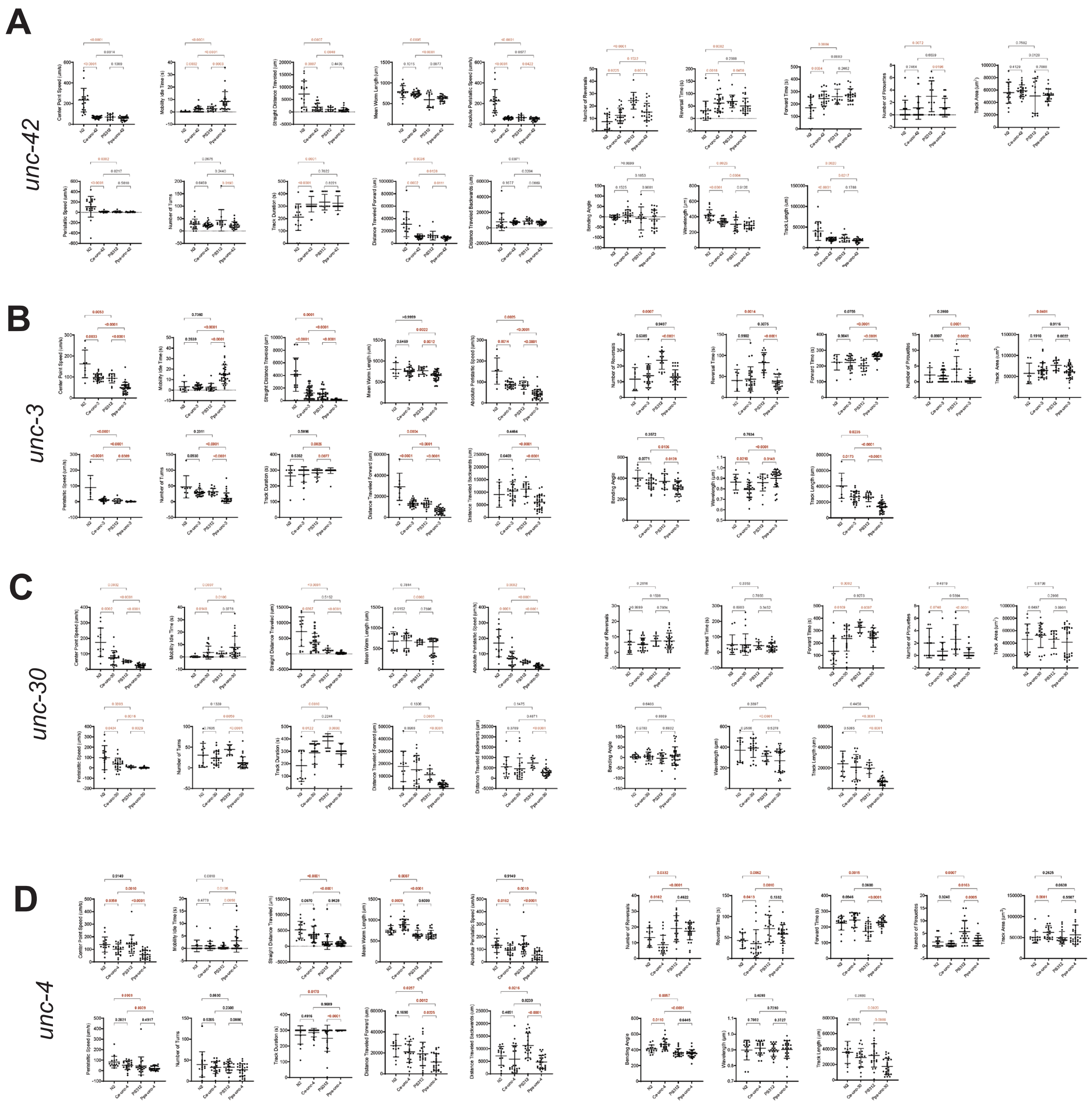


**Figure S9. Locomotion behavioral analysis of both *C. elegans* and *P. pacificus* wild-type and mutant strains for genes *unc-42, unc-3, unc-30* and *unc-4.*** **9. Locomotion behavioral analysis of both *C. elegans* and *P. pacificus* wild-type and mutant strains for genes *unc-42, unc-3, unc-30* and *unc-4.*** Five minute videos of freely moving worms were acquired using MultiWorm Tracker by MBF Biosciences. Automated tracking and editing were carried out with MBF software WormLab (*120*). **(A)** Quantitative analysis of tracking videos of *Ppa-unc-42* mutant worms. **(B)** *Ppa-unc-3* mutants. **(C)** *Ppa-unc-30* mutants. **(D)** *Ppa-unc-4* mutants.

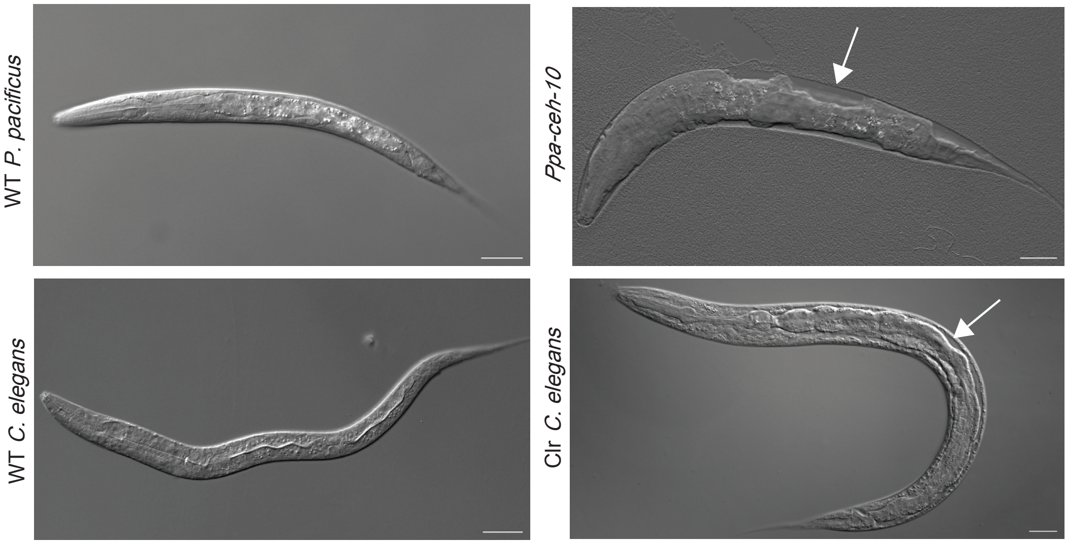


**Figure S10. Analysis of *Ppa-ceh-10* null mutation shows conservation of Clr phenotype and CAN molecular identity.** DIC images of a wild-type adult worm (top) and larvae containing a *Ppa-ceh-10* null mutation (bottom two panels). Arrows in images point to 'clear' (Clr) region in the body. The Clr phenotype in *Ppa-ceh-10* mutants is comparable to the Clr phenotype seen in a *C. elegans clr* mutant.

#### SUPPLEMENTARY TABLE LEGENDS

**Table S1. Conservation of homeobox proteins and other identity regulators in *P. pacificus* (Ppa) compared to *C. elegans* (Cel).** **Sheet 1:** Ppa orthologs of all non-Caenorhabditis-specific Cel homeoboxes. BLAST scores, e-values, % identities and % positives are results of Cel proteins BLASTed against the Ppa genome. **Sheet 2:** Ppa orthologs of non-homeodomain Cel terminal selectors. BLAST scores, e-values, % identities and % positives are results of Cel proteins BLASTed against the Ppa genome. **Sheet 3:** Ppa orthologs of Cel subtype selectors. BLAST scores, e-values, % identities and % positives are results of Cel proteins BLASTed against the Ppa genome. **Sheet 4:** Additional Ppa homeodomain proteins with no clear Cel ortholog (possible divergent homeodomain proteins). BLAST scores, e-values, % identities and % positives are results of closest Cel proteins BLASTed against the Ppa genome.

**Table S2. Terminal selector duplications in *P. pacificus*.** Ppa orthologs of Cel genes that have been found to have multiple copies that do not exist in the Cel genome. *mls-2*, *elt-1*, *ceh-31*, and *sox-2* all contain at least 1 duplicate in the *P. pacificus* genome. Expression patterns are either split (i.e. *mls-2*) between the duplicates or only one of the duplicates will have the expected expression pattern (i.e. *sox-2*).

**Table S3. Summary of molecular mutant phenotypes in *Ppa-unc-86*, *Ppa-unc-42*, *Ppa-unc-3*, *Ppa-unc-4*, and *Ppa-unc-30* null mutants.** Bolded and highlighted phenotypes are different from *C. elegans*.

**Table S4. Strains and CRISPR/Cas9 reagents.** Strains, as well as sequences of guides, repair templates, and alleles in this study.

**Table S5. HCR RNA-FISH reagents.** List of genes, amplifiers and fluorophores used in this study.
